# Repurposing Vanoxerine as a new antimycobacterial drug and its impact on the mycobacterial membrane

**DOI:** 10.1101/2022.11.29.517118

**Authors:** Alexander D. H. Kingdon, Asti-Rochelle Meosa John, Sarah M. Batt, Gurdyal S. Besra

**Author notes:** **Corresponding Author:** Alexander Kingdon.

## Abstract

*Mycobacterium tuberculosis* is a deadly pathogen, currently the leading cause of death worldwide from a single infectious agent through tuberculosis infections. If the End TB 2030 strategy is to be achieved, additional drugs need to be identified and made available to supplement the current treatment regimen. In addition, drug resistance is a growing issue, leading to significantly lower treatment success rates, necessitating further drug development. Vanoxerine (GBR12909), a dopamine re-uptake inhibitor, was recently identified as having anti-mycobacterial activity. Repurposing vanoxerine or its analogues to treat tuberculosis infections may allow a faster route to clinical use than novel drug discovery. However, its effects on Mycobacteria were not well characterised. Herein, we report vanoxerine as a disruptor of the membrane potential, inhibiting mycobacterial efflux and survival, with an undetectable level of resistance. This study suggests a mechanism of action for vanoxerine, which will allow for its continued development and optimisation for pre-clinical testing.

## Introduction

Tuberculosis is a major cause of death worldwide, accounting for approximately 1.6 million deaths in 2020 (WHO, 2022a). *Mycobacterium tuberculosis*, the causative agent, is a slow growing pathogenic species responsible for this deadly lung infection in over 10 million individuals every year (WHO, 2022a). Drug resistance to the current treatment regimen is a growing issue, representing 3% of new cases and 17% of re-infections (WHO, 2021). The treatment success rate against multi-drug resistant (MDR) tuberculosis is only 60% globally, necessitating new treatment options to combat this pandemic (WHO, 2022a; Pai et al., 2016).

For treatment of drug susceptible tuberculosis, a combination of four front-line drugs, ethambutol, isoniazid, pyrazinamide, and rifampicin, are taken for two months. Followed by isoniazid and rifampicin for a further four months (Nahid et al., 2016; Zumla et al., 2013). The combination therapy presents both financial and health burdens to patients, so any shortening or simplification of this treatment would provide a direct benefit to millions of patients every year. In addition, the treatment of MDR tuberculosis is more complicated, only recently being shortened to six-months with three to four drugs, from six drugs for up to 24 months (Nahid et al., 2016; Zumla et al., 2013; WHO, 2022b; Conradie et al., 2020).

In recent years, there has been some progress in the development of novel anti-tuberculosis drugs, with bedaquiline, delamanid and pretomanid all being approved for use (Zumla et al., 2013; Andries et al., 2005; Matsumoto et al., 2006; Skripconoka et al., 2012; Stover et al., 2000). However, all three drugs are restricted for use against MDR tuberculosis (Skripconoka et al., 2012; Centers for Disease Control and Prevention, 2013), and so do not allow for the shortening or simplification of the front-line drug regimen. In addition, some resistance has already been identified against these new drug compounds, necessitating further drug development (Andries et al., 2014; Hartkoorn et al., 2014).

One drug development approach which has been undertaken previously is drug repurposing (Corsello et al., 2017; Huang et al., 2011; Maitra et al., 2016), taking drugs with known activity and using their anti-mycobacterial properties for treatment of tuberculosis. Repurposing of drugs has been used for treating MDR tuberculosis in the past, including the use of linezolid, clofazimine and fluoroquinolones (Zumla et al., 2013). Currently, one-quarter of all ongoing clinical trials, for drug validation against tuberculosis, are focussed on repurposed drugs (WHO, 2022a). However, these drugs have all been known antibacterial compounds (Zumla et al., 2013). A recent study focussed on screening the Prestwick library, which consists of approved drugs against a large range of clinical implications, for activity against Mycobacteria (Kanvatirth et al., 2019). One of the drugs identified was vanoxerine (GBR12909), which showed activity against *Mycobacteria smegmatis, Mycobacteria bovis* BCG and *M tuberculosis* H37Rv (Kanvatirth et al., 2019).

Vanoxerine has been tested in several clinical trials, for two different clinical applications and has been well-tolerated by healthy volunteers (Søgaard et al., 1990; Preti, 2000; Lacerda et al., 2010; Laguna Pharmaceuticals, Inc., 2015). It has been identified as a dopamine re-uptake inhibitor (Lewis et al., 1999; Schmitt et al., 2008), for treatment of cocaine dependency, passing a phase I clinical trial. Vanoxerine has also been tested for use as an antiarrhythmic drug and passed a phase II clinical trial (Lacerda et al., 2010; Laguna Pharmaceuticals, Inc., 2015). It’s activity in this context was blocking the hERG potassium channel, which had no adverse effects on healthy volunteers but allowed treatment of atrial fibrillation (Lacerda et al., 2010; Cakulev et al., 2011; Obejero-Paz et al., 2015). However, the follow-up Phase III clinical trials was stopped due to adverse effects on the heart (ventricular proarrhythmia) in the treatment group (Piccini et al., 2016). These issues specifically occurred in patients with structural heart disease, which represented two thirds of the patients enrolled on the trial, hence, it was terminated early. If it is to be repurposed successfully, further trials are required to assess its safety against other patient groups (Piccini et al., 2016; Geng et al., 2020). Due to the knowledge of the human targets of vanoxerine, any future applications could monitor these targets for side effects. Future development of vanoxerine would also involve searching for analogues which retain their antimycobacterial effects but have reduced impact on dopamine reuptake.

Following the initial discovery of vanoxerine’s antimycobacterial properties (Kanvatirth et al., 2019), further characterisation of this drug’s effects on Mycobacteria was required. Herein, we have studied the impact of vanoxerine on mycobacteria, both phenotypic effects and transcriptomic impacts. Work has been undertaken with the aim of deconvoluting the target of vanoxerine and we suggest it targets the mycobacterial membrane, causing loss of the membrane electric potential and preventing efflux. In future, the presented work may allow the repurposing of vanoxerine or its analogues for treatment of tuberculosis.

## Methods

### General methods

Bacterial strains (outlined in Supplementary Table 1) were grown in Terrific broth (E. coli), BHI broth (C. glutamicum, *E. faecium*), LB broth (*A. baumanii, K. pneumoniae, P. aeruginosa, S. aureus*), 7H9 Middlebrook broth + 0.05% Tween-80 (M. smegmatis) or 7H9 Middlebrook broth + 0.05% Tween-80 + OADC (Oleic Acid Albumin Dextrose Complex, M. bovis BCG), unless stated otherwise. Bacterial cultures were grown at 37 °C and 180 rpm (or static for BCG) until mid-log (OD_600_ = 0.5-1.0), kanamycin (50 μg/ml) was added when required.

### Minimum inhibitory concentration (MIC) testing

Bacterial cultures were diluted to a final cell OD_600_ 0.05 in growth media. For pTIC protein over-expression, anhydrotetracycline (ATc, 200 ng/μl) was added. An intermediate compound plate was made using vanoxerine, performing a 1/2 or 2/3-fold serial dilution into dimethyl sulfoxide (DMSO). Each drug concentration (1 μl) was transferred into a 96-well polystyrene plate, followed by addition of diluted cell culture (99 μl). The plates were wrapped in foil and incubated (37 °C, 21 hours, or 6 days for BCG). Resazurin (30 μl, 0.02%, Acros Organics) was added to each well and the plate re-incubated (37 °C, 1 hour, 3 hours (M. smegmatis), or 24 hours (M. bovis BCG)). The resazurin fluorescence (excitation – 544 nm, emission - 590 nm) was measured in a BMG Labtech POLARstar Omega plate reader, normalised and cell survival calculated. For *A. baumanii, C. glutamicum, E faecium, K. pneumoniae, P. aeruginosa* and *S. aureus*, the OD_600_ was measured following 24-hours incubation, to calculate the percentage growth. For Checkerboard MICs, each well contained two drug compounds or DMSO, for a maximum 2% final concentration of DMSO. The fractional inhibitory concentration (FIC) was calculated (MIC_99_A in combination/MIC_99_A alone + MIC_99_B in combination/MIC_99_B alone) = FIC. If the FIC < 0.5, this represents synergy, 0.5 < FIC < 4, is indifference and FIC > 4, is antagonism (Odds, 2003).

### Mt-AroB protein expression and purification

The *M. tuberculosis aroB* gene was cloned into the pET28a vector using HiFi cloning. Briefly, the *aroB* gene was amplified from *M. tuberculosis* genomic DNA, using PCR to add on the complementary pET28a regions (forward primer - GCAGCCATCATCATCATCATATGACCGATATCGGCGCAC and reverse primer - GGCACCAGGCCGCTGCTGTGTCATGGGGCGCAAACTCC), while the pET28a vector was amplified to linearise at the point of insertion (forward primer - ATGATGATGATGATGGCTGCTGCCC and reverse primer - CACAGCAGCGGCCTGGTG). The NEBuilder^®^ HiFi DNA Assembly Cloning Kit was used, with a 1:2 ratio of vector to insert, and the reaction product transformed into *E. coli* XL10 competent cells, selecting on LB agar with kanamycin (50 μg/ml). The gene insertion into pET28a was confirmed by Sanger sequencing and the plasmid transformed into the *E. coli* BL21 (DE3) expression strain. An overnight culture (10 ml) of *E. coli* BL21 (DE3) was used to inoculate Terrific broth (1 litre) with kanamycin (50 μg/ml). The culture was grown (37 °C, 180 rpm) until mid-log growth was reached. Then IPTG (1 mM) was added and grown overnight (25 °C, 180 rpm). The cells were harvested by centrifugation (6,895 g, 10 minutes). The pelleted cells were resuspended in 500 mM NaCl, 30 mM Imidazole, 50 mM Tris-HCl buffer pH 7.8 (25 ml). The cells were lysed via sonication (3 minutes) and then centrifuged (31,360 g, 20 minutes). The clarified lysate was filtered (0.22 μM) and then loaded into a gravity-flow column containing HisPur Ni-NTA Resin (5 ml). The resin was washed with 500 mM NaCl, 30 mM Imidazole, 50 mM Tris-HCl buffer pH 7.8, before elution by increasing the imidazole concentration stepwise to 400 mM. The fractions containing MtAroB were pooled and dialysed overnight into 200 mM NaCl, 50 mM Tris-HCl buffer pH7.8. The protein was concentrated and flash frozen for storage at -80 °C.

### Fluorimetry assay for Mt-AroB compound binding

Mt-AroB protein (3 μL, 4.6 mg/ml) was added to 200 mM NaCl, 50 mM Tris-HCl buffer pH7.8 (400 μL) in a 1 cm quartz cuvette. This was equilibrated for 10 minutes (25 °C), before tryptophan fluorescence (excitation – 280 nm, emission – 340 nm) measurements were undertaken. Compounds of interest were added (1 μl per minute, 1 mM) and fluorescence was measured. The fluorescence changes associated with the buffer or DMSO (10%), were subtracted from the report values.

### Resistant mutant generation

Agar plates (5 ml, 7H11 + OADC) containing 2x, 2.5x, 5x and 10x MIC of vanoxerine were each inoculated with 1×10^8^ cells of *M. bovis* BCG. The plates were then incubated at 37 °C until the plates dried out, approximately 3 months.

### Ethidium bromide assays (accumulation and efflux)

A 96-well plate was prepared containing 1 μl of either DMSO, verapamil or vanoxerine, for final concentrations of 1%, 25 ug/ml or 3.3-52 μg/ml respectively. Mycobacterial cultures were washed and resuspended in PBS (4 ml, OD_600_ = 0.8) with tween-80 (0.05%) and ethidium bromide (0.625 μg/ml). For accumulation, glucose (0.4% (w/v) was added, and the diluted cell culture was immediately added to the 96-well plate. For efflux, verapamil (50 ug/ml) was added and the diluted culture was then incubated for 1 hour at 37 °C. The ethidium bromide loaded cells were centrifuged (3000 g, 8 min, 4 °C), and resuspended in cold PBS (4 ml, OD_600_ = 0.8) with tween-80 (0.05%). The cells were either added directly to the 96-well plate or supplemented with glucose (final concentration 0.4% (w/v)) before addition. In both assays, the fluorescence was then measured, in a BMG Labtech POLARstar Omega plate reader, every 60 seconds for 1 hour (emission – 544 nm, excitation – 590 nm), at 37 °C. Method adapted from (Rodrigues et al., 2021).

### DiOC_2_(3) Membrane potential assay

Mycobacteria cultures were washed and resuspended in PBS (3 ml, OD_600_ = 0.5) with tween-80 (0.05%) and DiOC_2_(3) (30 μM). The diluted culture was incubated (37 °C, 2 hours), before being dispensed (99 μl) into a 96-well plate. The fluorescence was measured every 90 seconds for 9 minutes in a BMG Labtech POLARstar Omega plate reader (excitation – 485 nm, emission – 520 nm, 620 nm), at 37 °C. The plate was then removed from the plate reader, 1 μl of either DMSO, CCCP, verapamil, vanoxerine or bedaquiline (final concentrations = 1%, 25 μM, 50 μg/ml, between 13-52 μg/ml or 0.5 μg/ml, respectively), was added to each well. The plate was then returned to the plate reader and readings continued for a further 50 minutes. Method adapted from (Chen et al., 2018; Li et al., 2019).

### Lipid extraction and thin-layer chromatography (TLC) analysis

*M. smegmatis* cultures were split into flasks (5 ml, OD_600_ = 0.3) containing either no drug, 13, 26 or 65 ug/ml vanoxerine, alongside C^14^ acetic acid (0.5 μCi per ml). These cultures were incubated at 37 °C for 24-hours, before cells were harvested by centrifugation (3,000 g, 10 minutes). The cell samples were stored at -20 °C until required. The pelleted cells were thawed and CH_3_Cl/MeOH/H_2_O (10:10:3, v/v/v, 2 ml) was added. The resuspension was incubated (2 hours, 50 °C), followed by addition of CH_3_Cl (1.75 ml) and H_2_O (0.75 ml). The lower organic phase was taken and washed twice with CH_3_Cl/MeOH/H_2_O (3:47:48, v/v/v, 2 ml). Then the lower organic phase was transferred to a new tube and dried. For TLC analysis, the lipids were resuspended in CH_3_Cl/MeOH (2:1, v/v, 200-500 μl) and spotted (5-20 μl) onto silica TLC plates corresponding to 10,000 counts. Thin layer chromatography was performed in CH_3_Cl/MeOH/H_2_O (80:20:2, v/v/v). The lipids were visualised by exposure of the TLC plate to an X-ray film for 1-week.

### RNA-sequencing

*M. bovis* BCG Pasteur was cultured to an OD_600_ 0.4 before exposure to DMSO, 15 or 30 ug/ml vanoxerine for eight hours in four biological replicates each. Cells (10^8^) were pelleted, flash frozen in liquid nitrogen and stored at −80 °C and then sent to Genewiz (Azenta Life Sciences) for RNA extraction and sequencing. The raw transcriptomic outputs were mapped onto the *M. bovis* BCG Pasteur (GCA_000009445.1) and *M. tuberculosis* H37Rv (GCA_000283295.1) genomes. Annotation and clustering of the significantly dysregulated gene transcripts was performed using the DAVID server (Sherman et al., 2022; Huang et al., 2009). The transcriptomic data has been submitted to the European Nucleotide Archive (ENA), accession number PRJEB57729.

## Results

### Vanoxerine showed selectivity for Mycobacterial species

Vanoxerine was initially identified to have activity against *M. smegmatis* and *M. bovis* BCG during a high-throughput screening of the Prestwick library (Kanvatirth et al., 2019). Further work confirmed it’s activity against *M. tuberculosis* H37Rv directly (Kanvatirth et al., 2019). This was the first time vanoxerine had been reported to have anti-bacterial activity. To determine its spectrum of activity, testing was undertaken on clinically relevant Gram-negative (*A. baumanii, K. pneumoniae, P. aeruginosa*) and Gram-positive (*E. faecium* and *S. aureus*) species, alongside testing *Corynebacterium glutamicum*, to determine the MIC (Table 1, Supplementary Figure 1). The MIC_99_ was found by calculating normalised percentage survival using DMSO and antibiotic controls, with the lowest drug concentration which caused a 1% or lower percentage survival being the MIC_99_. The selectivity of vanoxerine for Mycobacteria was highlighted, with low MIC_99_s for *M. tuberculosis and M. smegmatis* of 14 and 26 μg/ml respectively, compared with greater than 52 μg/ml for *S. aureus* and *P. aeruginosa*. Vanoxerine exhibited no activity against Gram-negative species (Table 1), including species with EDTA-permeabilised outer membranes (data not shown), suggesting no equivalent target is present. Amongst Gram-positive species, *Enterococcus faecium* was inhibited by the drug, but no effect was found against *Staphylococcus aureus*, indicating some specificity amongst Gram-positive species. While the liquid MIC_99_ for *M. bovis* BCG was much higher than *M. tuberculosis*, the solid agar MIC_99_ was determined to be 15.2 μg/ml.

**Table 1:** Minimum Inhibitory Concentrations (MICs) Against Representative Species.

| Species | MIC<sub>99</sub> (µg/ml) |
| --- | --- |
| *M. smegmatis* | 23-35 |
| *M. bovis* BCG | 26 |
| *M. tuberculosis* H37Rv | 14 |
| *C. glutamicum* | 15.5 |
| *E. faecium* 64/3 | 23 |
| *S. aureus* | >52 |
| *A. baumannii* | >52 |
| *K. pneumoniae* | >52 |
| *P. aeruginosa* | >52 |

### AroB is not the mycobacterial target of Vanoxerine

In addition to reporting the anti-mycobacterial activity of vanoxerine, previous work indicated a potential target, AroB, a 3-dehydroquinate synthase enzyme involved in the shikimate pathway for aromatic amino acid synthesis (Kanvatirth et al., 2019; Dutra De Mendonça et al., 2007). To confirm a vanoxerine-AroB interaction, we re-cloned the *M. tuberculosis aroB* gene into the same pVV16 expression vector used for constitutive overexpression and additionally, into the pTIC6a vector, used for anhydrotetracycline (ATc) inducible expression, both in *M. smegmatis*. These plasmids were transformed in *M. smegmatis*, and vanoxerine percentage survival curves repeated, comparing empty vector controls to the AroB overexpression plasmids (Figure 1A). In contrast to the previous work (Kanvatirth et al., 2019), there was no difference between the empty vector and AroB over-expression conditions. For the pTIC6a strains, the MIC of both empty and *aroB* vectors may have shifted due to presence of ATc at 200 ng/μL used to induce overexpression (Figure 1A).

**Figure 1:**
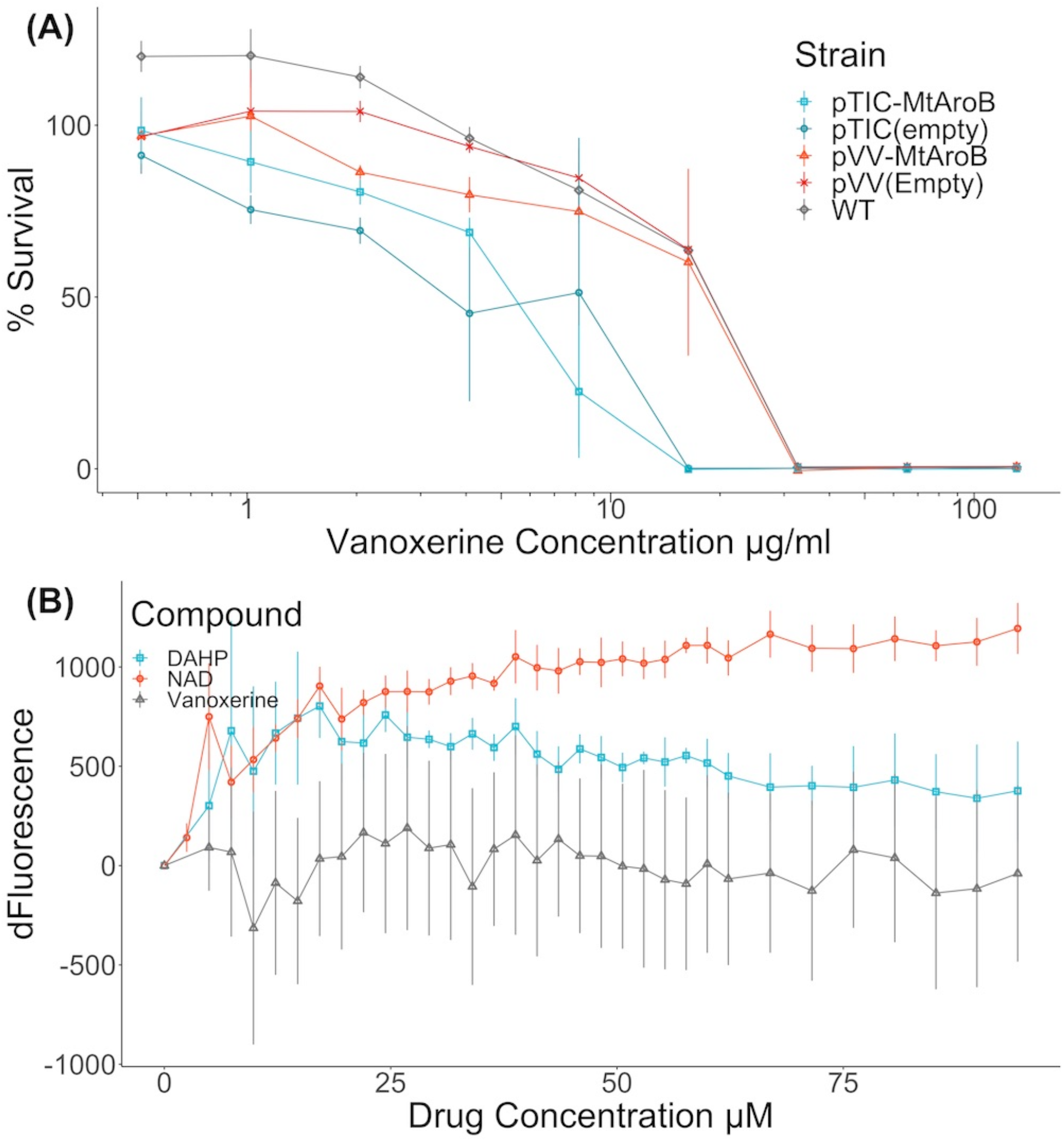
AroB overexpression had no effect on vanoxerine killing and vanoxerine does not bind to the purified protein. **(A) % Survival Curve of *M. smegmatis* over-expressing *M. tuberculosis* AroB relative to empty vector controls, against Vanoxerine.** All *M. smegmatis* strains were grown in the presence of vanoxerine, with resazurin used to quantify survival. The % survival was calculated compared to DMSO and rifampicin controls. N=3 (**B) Fluorimetry using the tryptophan fluorescence of purified *M. tuberculosis* AroB protein titrated against Vanoxerine and known binding compounds**. The change in fluorescence (dFluorescence) was calculated compared to the baseline fluorescence, following each drug addition, with the changes due to solvent (buffer or 10% DMSO) subtracted. DAHP (substrate) = 3-deoxy-D-arabinoheptulosonate 7-phosphate, NAD (co-factor) = nicotinamide adenine dinucleotide. N=2

Due to the ambiguity of these contrasting results, we expressed and purified the *M. tuberculosis* AroB protein (Supplementary Figure 2), to allow direct binding interactions to be studied. Following protein purification, tryptophan fluorescence was used to assess the binding of compounds to the AroB protein. In addition to vanoxerine, the binding of 3-deoxy-D-arabinoheptulosonate 7-phosphate (DAHP), the substrate of AroB and nicotinamide adenine dinucleotide (NAD), the co-factor of AroB (Dutra De Mendonça et al., 2007, 2011), were utilised as controls. Both DAHP and NAD caused large shifts in tryptophan fluorescence during addition (Figure 1B), assumed to represent protein conformational changes in response to compound binding. However, upon addition of vanoxerine, no equivalent change in fluorescence occurred (Figure 1B). Overall, these results suggested AroB is not the target of vanoxerine, and hence additional work was undertaken to identify vanoxerine’s mechanism of action.

### No resistance could be generated against Vanoxerine in *M. bovis* BCG

In an effort to find the true Mycobacterial target of vanoxerine, spontaneous resistant mutant generation was attempted in *M. bovis* BCG (Abrahams and Besra, 2020). However, no resistant mutants could be generated, with no growth occurring at even 2x MIC_99_ of vanoxerine (30 μg/ml). This suggested the rate of resistant generation is below 10^8^. In addition, an *M. bovis* BCG strain with a *recG* mutation was used, due to its increased mutation rate (Ley et al., 2019; Batt et al., 2015). However, the Δ*recG* strain also failed to acquire any spontaneous resistant mutations, suggesting a rate of resistance even lower than 10^8^. This is promising for the future use of vanoxerine; however, this approach did not allow a target to be identified.

### Vanoxerine inhibits ethidium bromide efflux

Many current antimycobacterial drugs target either the cell envelope or cellular energetics (Andries et al., 2005, p.203; Pethe et al., 2013; Zumla et al., 2013). Hence, this led us to investigate the impact of vanoxerine on the cell envelope, to study the impact on membrane integrity and energy requiring processes such as efflux. Ethidium bromide uptake and efflux in Mycobacteria has previously been described (Rodrigues et al., 2021) and allows fluorescent monitoring of the impact of drugs on membrane permeability and efflux. The MIC_99_ for ethidium bromide against both *M. smegmatis* and *M. bovis* BCG was determined to be 6.25 μg/ml. For the assays, 1/10th MIC_99_ of ethidium bromide was used, so the cell viability was not affected during the assay. In addition, glucose was added to provide an energy source, enabling the cells to efflux more ethidium bromide.

During the 60-minute assay window, vanoxerine caused a significant difference in the accumulation of ethidium bromide inside both *M. smegmatis* and *M. bovis* BCG (Figure 2A, Supplementary Figure 3A). Vanoxerine concentrations as low as 0.125x MIC_99_ (3.27 μg/ml) led to more ethidium bromide accumulation than the DMSO control. The effect was also concentration dependent, with 2x MIC_99_ (52 μg/ml) of vanoxerine causing 2.6x more fluorescence than DMSO and 1.2x more fluorescence than 0.5x MIC_99_ (13 μg/ml) across the 60-minutes, due to ethidium bromide accumulation. As there could be several explanations for these results, including membrane disruption/lysis, efflux inhibition or cell energetic effects, further work was undertaken.

**Figure 2:**
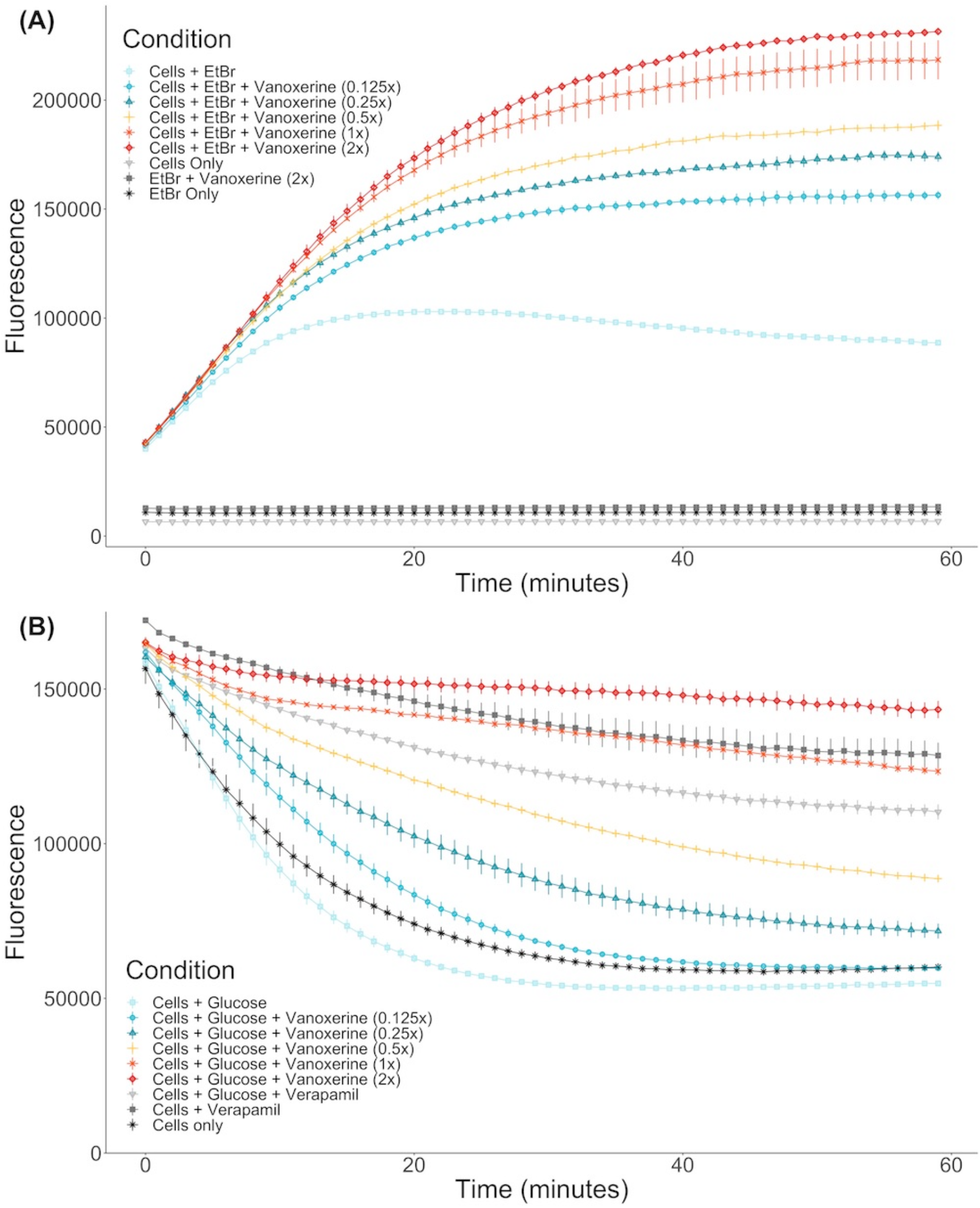
Vanoxerine inhibits the efflux of ethidium bromide from *M. smegmatis*. **(A) Vanoxerine treatment leads to accumulation of ethidium bromide in *M. smegmatis* at sub-inhibitory concentrations.** *M. smegmatis* (OD_600_ = 0.8) was re-suspended in PBS + 0.05% tween-80 + 0.4% glucose + 0.625 μg/ml ethidium bromide. The cells were immediately added to a 96-well plate containing varying vanoxerine concentrations and the fluorescence was monitored. MIC = 26 μg/ml. N=3. **(B) Vanoxerine prevents the efflux of ethidium bromide in *M. smegmatis***. Verapamil was used to cause ethidium bromide accumulation in *M. smegmatis* for 1 hour. Then the cells were resuspended in fresh PBS and either added directly, or mixed with glucose to 0.4% before adding, to a 96-well plate containing DMSO, vanoxerine or verapamil. The fluorescence was monitored to look at efflux of ethidium bromide. MIC=26 μg/ml. N=3.

To study inhibition of efflux by vanoxerine, Mycobacterial cells were allowed to accumulate ethidium bromide in the presence of a non-toxic efflux inhibitor verapamil. The verapamil MIC_99_ against *M. smegmatis* and *M. bovis* BCG was greater than 100 μg/ml, having limited impact on metabolic activity at the highest tested concentration. Following ethidium bromide accumulation in Mycobacteria, verapamil was removed.

The efflux could then be monitored as loss of fluorescence. In the absence of vanoxerine or verapamil, a clear decrease in fluorescence could be observed, approximately 65% (Figure 2B). In the presence of 50 μg/ml verapamil, the decrease in fluorescence was reduced to approximately 32%, indicating inhibition of ethidium bromide efflux from the cell. In the presence of 1x and 2x MIC of vanoxerine (26 or 52 μg/ml), the drop in fluorescence was further reduced to 25% and 13% respectively (Figure 2B). This indicated efflux was being inhibited by vanoxerine. Normal rates of efflux (63% drop in fluorescence) were only evident at vanoxerine concentrations of 0.125x MIC (3.27 μg/ml) and lower. A similar impact on efflux was also observed in *M. bovis* BCG, but with lower rates of accumulation and efflux (Supplementary Figure 3B).

### Vanoxerine impacted the membrane potential (ΔΨ)

As vanoxerine was able to inhibit efflux of ethidium bromide, the voltage sensitive dye DiOC_2_(3) was used to monitor the electric potential (ΔΨ) of the membrane in response to vanoxerine treatment, as ΔΨ disruption would prevent the majority of efflux (Remm et al., 2022). DiOC_2_(3) partitions across phospholipid membranes proportionally to the ΔΨ present (Chen et al., 2018; Chawla et al., 2012; Hudson et al., 2020; Li et al., 2019; Novo et al., 1999), fluorescing red inside cells and green in solution. It was investigated whether vanoxerine disrupts the electric potential of the membrane by measuring DiOC_2_(3) membrane partitioning. Hence, *M. bovis* BCG was pre-incubated with DiOC_2_(3) for two hours prior to drug addition, to allow the dye to equilibrate across the membrane proportional to the ΔΨ. The red/green fluorescence ratio fluctuated around 1.75 in the absence of compound, reducing to 1.60 across the 60-minute assay. Vanoxerine was able to disrupt the dye partitioning in a concentration dependent manner, with 52 μg/ml vanoxerine reducing the red/green fluorescence to 1.09 after 50 minutes, suggesting disruption of the ΔΨ (Figure 3). The membrane potential was still disrupted at sub-MIC amounts of vanoxerine, with 13 μg/ml (0.5x MIC) leading to a red/green fluorescent ratio of 1.30. CCCP is a protonophore which is known the disrupt both the ΔΨ and proton gradient (ΔpH) of the membrane (Chen et al., 2018) and had the lowest red/green fluorescent ratio in this assay, of 0.95. Vanoxerine disrupted the DiOC_2_(3) dye partitioning at a slower rate compared to CCCP. The assay was specific to the ΔΨ disruption, rather than general proton motive force (PMF) disruption. Bedaquiline confirmed the specificity as it is known to only disrupt the ΔpH of the PMF, and hence had a response similar to DMSO in this assay, a red/green fluorescent ratio of 1.63 (Feng et al., 2015). In addition, kanamycin had no impact on DiOC_2_(3) partitioning, with a red/green fluorescent ratio of 1.63 after 50 minutes of incubation, suggesting a general bactericidal response is less likely to explain the dyes disruption.

**Figure 3:**
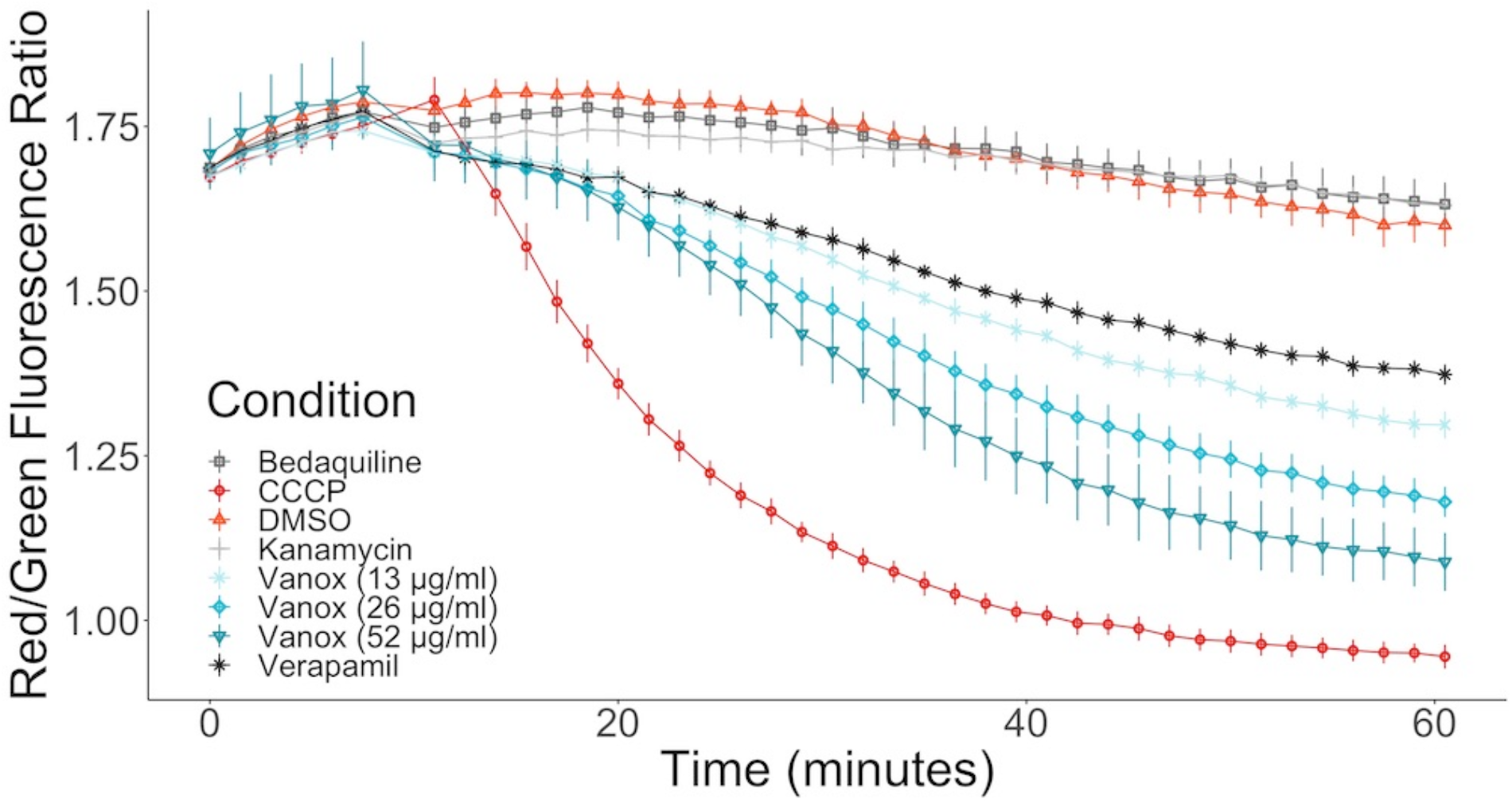
Vanoxerine disrupts the membrane potential of *M. bovis* BCG, disrupting already partitioned DiOC_2_(3) dye. Bedaquiline = 0.5 μg/ml, CCCP = 25 μM, DMSO = 1%, Kanamycin = 100 μM, Verapamil = 50 μg/ml. *M. bovis* BCG was incubated with 30 μM DiOC_2_(3) for 2 hours to allow dye partitioning across the membrane, the fluorescence was then measured for 10 minutes. Then either vanoxerine or controls were added, and the fluorescence was measured for a further 50 minutes. N=3.

### Vanoxerine may potentiate the activity of other anti-mycobacterial drugs

As vanoxerine appears to inhibit efflux from Mycobacteria, it was hypothesised that any anti-mycobacterial drugs which are known to be pumped out by the cell, may have increased efficacy if used alongside vanoxerine. Resistance to bedaquiline, clofazimine and FNDR-20081 have been shown to occur via mutations in Rv0678, the transcriptional repressor of the *mmpL5-mmpS5* operon, leading to up-regulation of MmpL5 (Hartkoorn et al., 2014; Andries et al., 2014; Remm et al., 2022; Kaur et al., 2021). This suggests that all three drug compounds are pumped out of the cell by MmpL5. Checkerboard MICs against *M. bovis* BCG were set up for bedaquiline, clofazimine and FNDR-20081 against vanoxerine. In addition, to test for synergy with front-line drugs, a rifampicin vs vanoxerine checkerboard was set up. In the presence of sub-inhibitory concentrations of vanoxerine (10.3 μg/ml), the MIC_99_ of rifampicin, clofazimine and FNDR-20081 were reduced by 2x (0.05 to 0.025 μg/ml, 4.8 to 2.4 μg/ml, and 24.5 to 12.3 μg/ml, respectively, Figure 4). The FIC values were calculated as 0.676 for rifampicin and 0.896 for clofazimine and FNDR-20081, with vanoxerine. These FIC values were all over 0.5 and so indicate no interactions were occurring; hence, further work needs to be undertaken to determine if the observed MIC_99_ reductions are due to the presence of the vanoxerine or due to experimental set-up.

**Figure 4:**
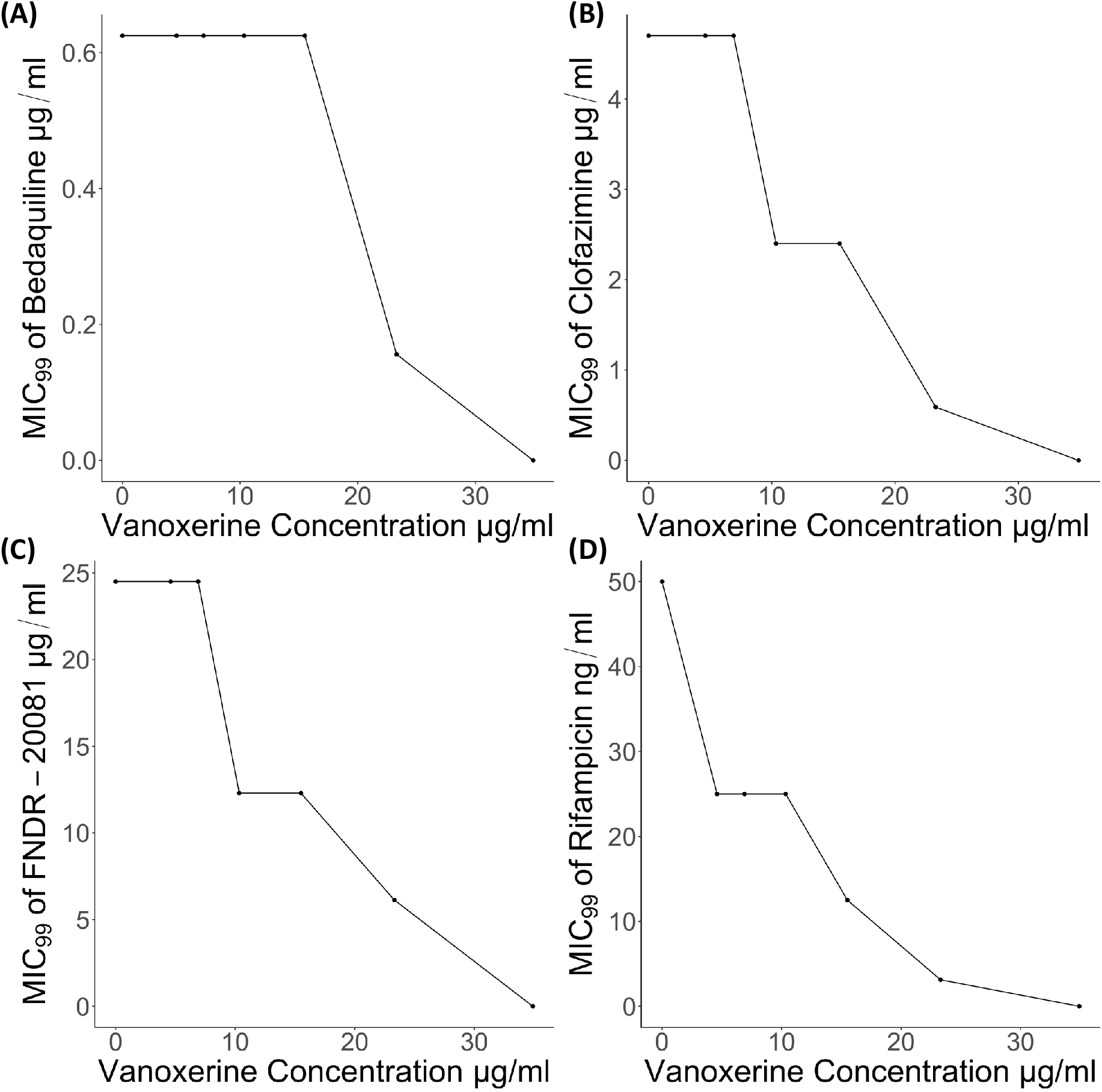
Vanoxerine potentiates the effects of other anti-mycobacterial drugs, including Clofazimine, Ethidium bromide, FNDR-20081 and Rifampicin. **(A) Bedaquiline. (B) Clofazimine. (C) FNDR-20081. (D) Rifampicin.** Resazurin turnover was used as a proxy for survival of *M. bovis* BCG following 7-days of drug treatment. The checkerboard MICs was created to assess seven vanoxerine concentrations against eleven anti-mycobacterial drug concentrations, with a final DMSO concentration of 2%. The MIC_99_ of each drug is plotted against the vanoxerine concentration in that row of the plate. These plots represent the average of three independent checkerboard plates. concentrations against eleven anti-mycobacterial drug concentrations, with a final DMSO concentration of 2%. The MIC_99_ of each drug is plotted against the vanoxerine concentration in that row of the plate. These plots represent the average of three independent checkerboard plates.

In contrast, the FIC value for bedaquiline was 1.13, also suggesting no interaction, but corresponded to no difference to the bedaquiline MIC_99_ in the presence of 10.3 μg/ml vanoxerine (Figure 4A).

### Vanoxerine induced clear transcriptomic changes in *M. bovis* BCG

As the inhibition of efflux and disruption of electric potential occurred at sub-inhibitory concentrations of vanoxerine (Figure 2B, Figure 3), this suggested that other pleotropic effects may be leading to cell death. Hence, transcriptomic analysis of *M. bovis* BCG was undertaken using RNA-sequencing following 8-hours of vanoxerine treatment. Two different drug concentrations were used, to investigate if any transcriptomic differences were concentration dependent. Vanoxerine induced clear transcriptomic differences compared to the DMSO only control, with clear separation between the no drug and 30 μg/ml vanoxerine (Figure 5A). 96% of the variance across the eight samples was due to the treatment conditions. The transcription of over 800 genes was significantly differentially regulated at 30 μg/ml of vanoxerine compared to a DMSO only control (fold change = 2 or more, pvalue <0.05, (Figure 4B)). At 15 μg/ml vanoxerine, 322 of these gene transcripts were still significantly dysregulated compared to DMSO (Supplementary Figure 4). Comparing between vanoxerine treatment conditions; there were 304 genes in common, that were significantly dysregulated. Only 17 significantly dysregulated gene transcripts were unique to 15 μg/ml vanoxerine, whilst 510 gene transcripts were uniquely dysregulated at 30 μg/ml vanoxerine. The list of 304 common genes could be narrowed down to 31 gene transcripts that were significantly dysregulated in a concentration dependent manner, when comparing 15 μg/ml vanoxerine as a baseline to 30 μg/ml vanoxerine (Supplementary Figure 5).

The significantly dysregulated genes at 30 μg/ml vanoxerine were functionally annotated and clustered using the DAVID server (Sherman et al., 2022; Huang et al., 2009), to identify pathways or biological processes that were either up-or down-regulated in response to vanoxerine (Figure 4C). In relation to up-regulated transcripts, these clustered to include: membrane stress responses; mycobactin synthesis; folate & riboflavin biosynthesis; oxidoreductase activity; polyketide synthesis; protein export; efflux; transporter proteins and drug de-toxification. In contrast, the down-regulated pathways include: oxidative phosphorylation; mycolic acid biosynthesis (Fas-I & FasII); cell wall biogenesis; amino acid biosynthesis; carbon metabolism and protein biosynthesis. The upregulation of gene transcripts involved in membrane stress, protein export, efflux and other transporters provide further evidence that vanoxerine’s mechanism of action inhibits these processes.

As only 31 genes were significantly dysregulated in a concentration dependent manner, these were investigated in more detail (Supplementary Table 2). There was little consensus of function among the significantly upregulated genes, with the majority being of unknown function. The efflux pump MmpL5 was significantly upregulated, which could be due to an increasing impact on efflux at higher vanoxerine concentrations. The majority of significantly down-regulated transcripts were part of the mycolic acid biosynthetic pathway. Explanations for this result include the high energetic costs associated with mycolic acid production, indirect inhibition of MmpL3, or vanoxerine directly impacting this pathway.

### Mycolic acid down-regulation occurred following vanoxerine treatment, but direct inhibition is unlikely

To gain further evidence for the impact on mycolic acid biosynthesis, in addition to the four genes which were transcriptionally downregulated in a concentration dependent manner, the effect of vanoxerine on all the genes in the pathway was investigated (Table 2). Treatment with vanoxerine led to transcriptional repression of mycolic acid biosynthesis, including all the enzymes in FAS-I and FAS-II. The only gene transcript which was not downregulated was *mmpL3*, although its regulation was not significantly different from the DMSO control, even using 30 μg/ml vanoxerine.

**Table 2:** Gene Expression of Mycolic acid synthesis pathway - comparing the DMSO control to 30 μg/ml MIC Vanoxerine.

| ID | log2FoldChange | pvalue | padj |
| --- | --- | --- | --- |
| ***accD6*** | -1.4342963 | 8.57E-125 | 2.24E-123 |
| ***acpM*** | -2.0530831 | 6.07E-23 | 2.27E-22 |
| ***cmrA (Rv2509)*** | -0.5538547 | 6.72E-26 | 2.76E-25 |
| ***fabD*** | -1.7924797 | 4.62E-79 | 6.13E-78 |
| ***fabG1*** | -1.7545573 | 1.31E-105 | 2.69E-104 |
| ***fabH*** | -0.394042 | 0.00055311 | 0.00082604 |
| ***fadD32*** | 0.11059707 | 0.37003607 | 0.4089094 |
| ***fas*** | -0.9664376 | 5.02E-52 | 4.06E-51 |
| ***fbpA*** | -1.402825 | 2.07E-77 | 2.66E-76 |
| ***fbpB*** | -2.3725639 | 2.69E-179 | 1.51E-177 |
| ***fbpC*** | -0.5545252 | 0.00031762 | 0.00048482 |
| ***hadA*** | -1.4822077 | 1.08E-07 | 2.07E-07 |
| ***hadB*** | -1.0323571 | 1.05E-19 | 3.50E-19 |
| ***inhA*** | -1.7253415 | 1.92E-48 | 1.44E-47 |
| ***kasA*** | -2.1172224 | 8.97E-74 | 1.09E-72 |
| ***kasB*** | -2.0727098 | 5.37E-139 | 1.78E-137 |
| ***mmpL3*** | 0.79100589 | 1.03E-32 | 5.21E-32 |
| ***pks13*** | -0.1798483 | 1.84E-07 | 3.47E-07 |

To investigate whether this transcriptional down-regulation translated into loss or reduction of mycolic acids *in vitro*; lipid extraction was undertaken on *M. smegmatis* treated with vanoxerine. The lipids were labelled using C^14^ acetic acid at the same time as drug treatment. The extracted lipids were separated using thin-layer chromatography (TLC) and visualised using X-ray film (Figure 5A). Loss of trehalose mono-mycolate (TMM) occurred at 0.5x MIC of vanoxerine and trehalose di-mycolate

**Figure 5:**
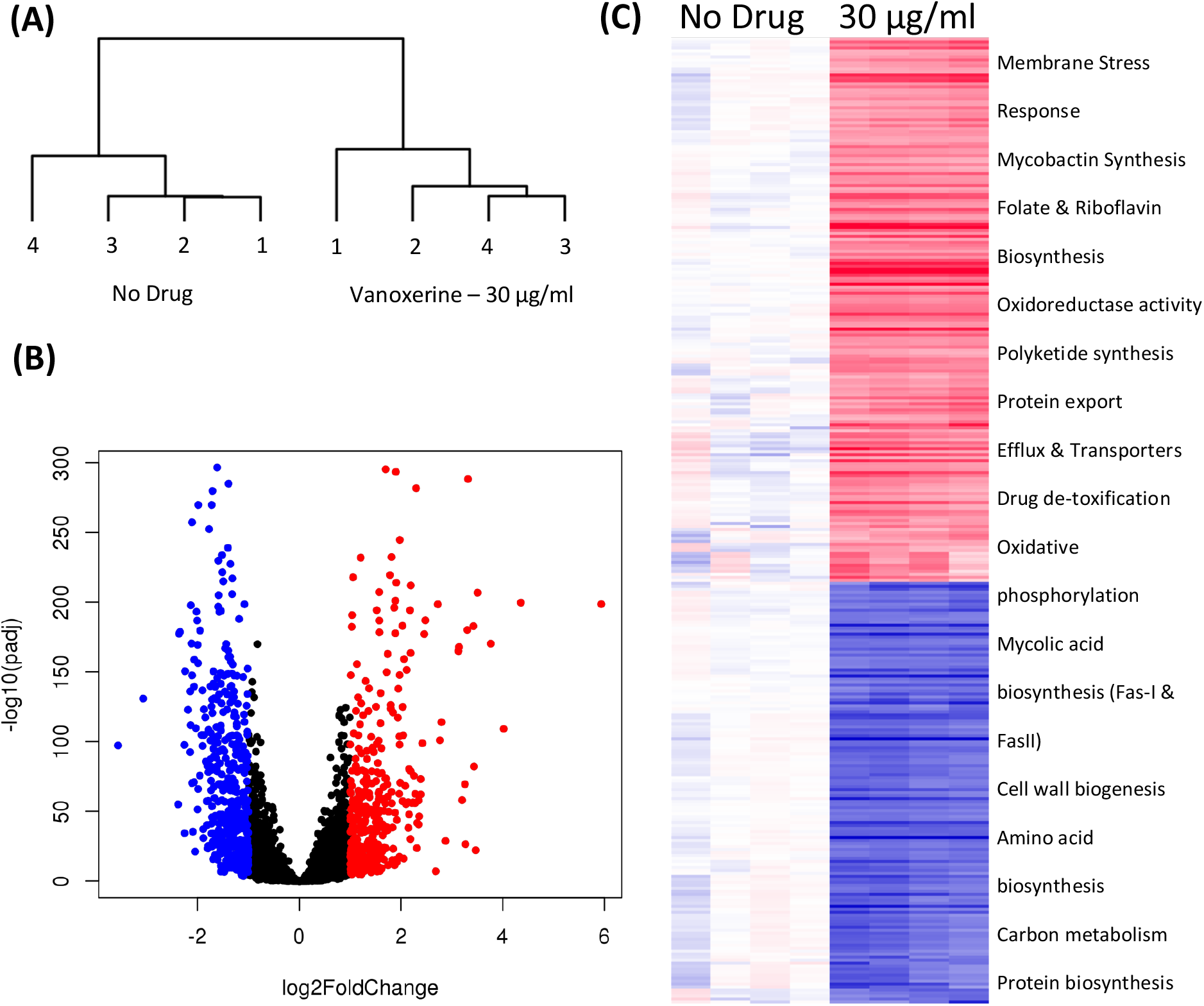
Transcriptomic changes following drug treatment of *M. bovis* BCG relative to a DMSO only control. **RNA sequencing was performed on four independent samples from each condition.** (A) A dendrogram, highlighting the sample differences present, due to transcriptional variation between the DMSO-only samples and 30 μg/ml vanoxerine treated samples, following 8-hours of drug treatment. (B) Volcano plot indicating the significantly dysregulated genes. Highlighted points have a fold change > 2 and a p-adjusted value of <0.05; red = up-regulated, blue = down-regulated. (C) Clustered comparison of transcriptional differences between DMSO-only and 30 μg/ml vanoxerine. Biological functions of the dysregulated genes are listed.

(TDM) at 2.5x MIC of vanoxerine. The loss of TMM was quantified by comparing the radioactive counts on the TLC plate in the presence or absence of vanoxerine (Figure 5B). A drop in TMM from 7% of the total counts to less than 1%, confirmed it is not just lower growth in the presence of vanoxerine which is caused this decrease. As the mycolic acids are essential to mycobacteria, this might be another mechanism of vanoxerine inhibition.

To investigate further, we chose to study vanoxerine’s impact on *C. glutamicum*, a species where the mycolic acids are not essential. As vanoxerine could inhibit the growth of *C. glutamicum*, with an MIC_99_ of 15.5 μg/ml (Table 1), comparable to the MIC_99_ of *M. tuberculosis*, suggesting mycolic acid biosynthesis is not the only target of vanoxerine. A *C. glutamicum* Δpks mutant, which does not synthesise mycolic acids, was compared to the wild-type strain for its survival following vanoxerine treatment (Figure 5C). No shift in % growth curve or difference in MIC_99_ was observed, suggesting mycolic acid biosynthesis inhibition is not the main mode of inhibition of vanoxerine. In addition, the over-expression of several genes in the mycolic acid biosynthetic pathway was undertaken in Mycobacteria (Supplementary Figure 6). However, no differences in survival to vanoxerine treatment were found, including for MmpL3 over-expression, providing further evidence that this is not a direct target of vanoxerine.

**Figure 6:**
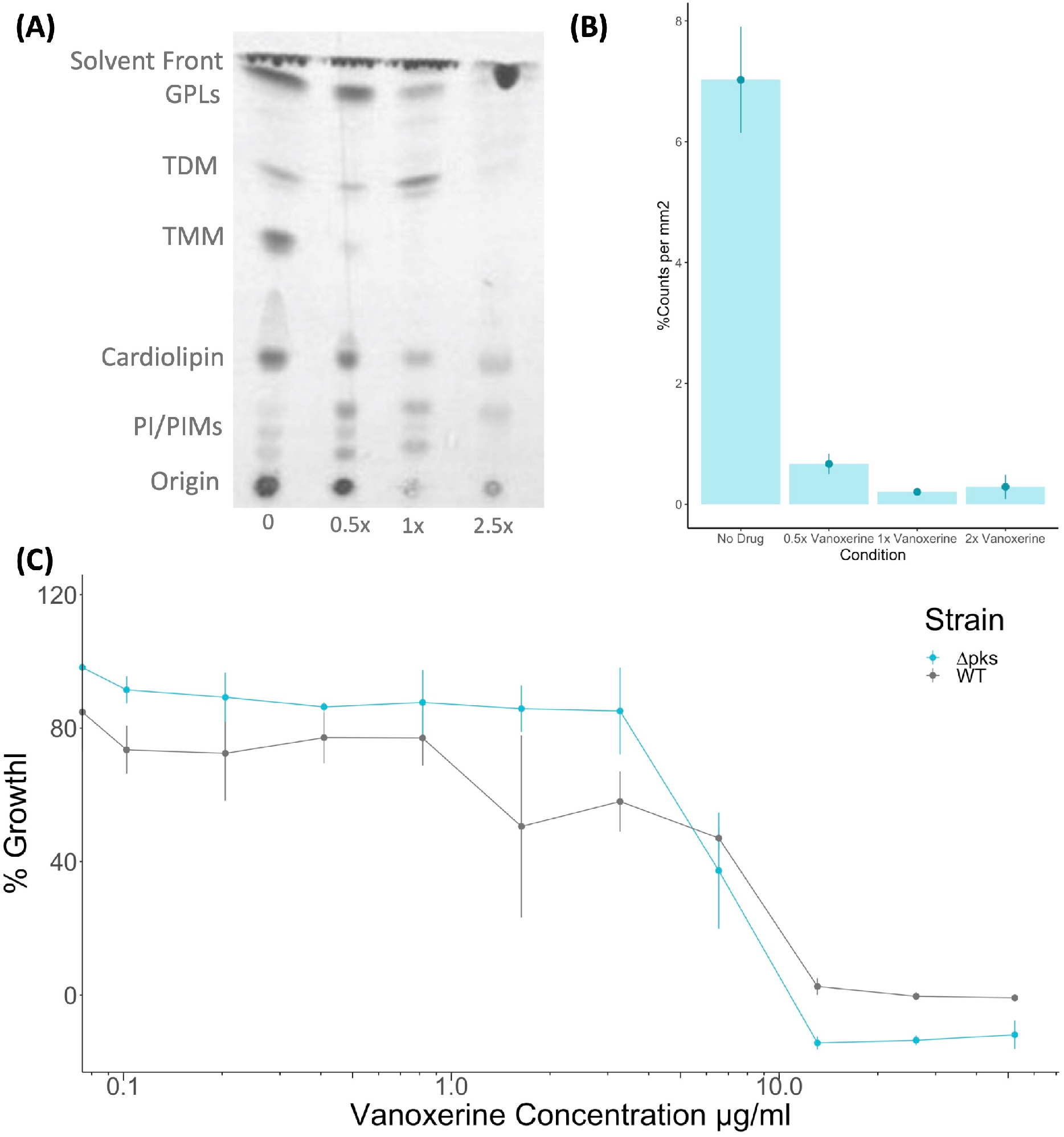
Vanoxerine treatment impacts the mycolic acids in *M. smegmatis*, but this inhibition is unlikely to be the main mechanism of inhibition. **(A) Thin Layer Chromatography of *M. smegmatis* lipid extractions, showing loss of TMM.** *M. smegmatis* wild-type was treated with vanoxerine for 24-hours, followed by a total lipid extraction. Thin layer chromatography was performed (CH_3_Cl/MeOH/H_2_O, 80:20:2, v/v/v), followed by exposure to an X-ray film to image. MIC = 26 μg/ml GPL = Glycopeptidolipids, TDM = Trehalose dimycolates, TMM = Trehalose monomycolates, PI = Phosphatidylinositol, PIMs = phosphatidylmannosides. Representative image from N=4. **(B) Densiometric analysis of TMM loss following vanoxerine treatment**. The same silica plates were exposed to a storage phosphor screen and then scanned to quantify the radioactivity of each spot. The counts for the TMM spot were compared to the total counts in each lane, to generate a %counts per mm^2^. N=4. **(C) %Growth comparing *C. glutamicum* WT vs Δpks13 mutant in the presence of vanoxerine**. *C. glutamicum* was incubated with vanoxerine for 24 hours, before OD_600_ was measured. The % survival was calculated compared to DMSO only and rifampicin controls. N=3.

## Discussion

This study has highlighted that vanoxerine has a limited spectrum of antibacterial activity, mainly targeting the Mycobacteriales, alongside some other Gram-positive species. Contrary to previous claims (Kanvatirth et al., 2019), AroB is unlikely to be the Mycobacterial target of vanoxerine. Rather, vanoxerine disrupts the membrane’s electric potential, causing downstream disruption of mycobacterial energetics. This finding has been supplemented with evidence of the inhibition of efflux and the disruption of transport of substances across the membrane. Finally, vanoxerine may have some ability to potentiate other anti-mycobacterial drugs, although further work is needed to confirm this effect.

The lack of AroB binding interactions *in vitro*, and the absence of a shift in the MIC following overexpression of the protein *in vivo*, are in direct contrast to the previous results (Kanvatirth et al., 2019). In addition, the position and mutations found in the *aroB* gene of the four ‘resistant mutant strains’ of *M. smegmatis* following vanoxerine treatment (Kanvatirth et al., 2019), do not affect the *M. tuberculosis* AroB homologue and so could not confer resistance. One mutation is in a terminal unstructured region not present in the *M. tuberculosis* AroB protein, and the other three *M. smegmatis* mutations converted the residues to the equivalent *M. tuberculosis* AroB protein residues (PDBE ID 3QBE) (Table 1). The evidence provided indicates vanoxerine does not interact with AroB and it is unlikely to be the target.

Previous attempts to generate resistant mutants to vanoxerine were performed in *M. smegmatis* (Kanvatirth et al., 2019), as a standard mode-of-action determination method used for anti-mycobacterial drugs, followed by whole-genome sequencing of the mutants (Abrahams and Besra, 2020). This approach was repeated in *M. bovis* BCG. However, during the course of this work, several attempts were made to generate resistant mutants to vanoxerine, but no mutants could be isolated, including use of a recG mutant (Batt et al., 2015). The lack of spontaneous resistance is promising for the longevity of the drug, due to the increasing levels of MDR tuberculosis (WHO, 2021), and resistance to the most recently approved anti-mycobacterial drugs, bedaquiline, delamanid, and pretomanid (Zumla et al., 2013). The lack of *in vivo* resistance may also indicate pleotropic effects on the cell, reducing resistance development.

The retention of ethidium bromide by Mycobacteria provides strong evidence that vanoxerine inhibits efflux. Cell lysis or pore formation are unlikely to be the mechanism of action, as a faster decrease in ethidium bromide fluorescence would be the expected outcome. Efflux inhibition could occur via several mechanisms, including inhibition of ATP synthesis, disruption of membrane energetics, or direct efflux pump inhibition (Remm et al., 2022). Direct efflux pump inhibition is less likely, due to several types of efflux pumps being involved in ethidium bromide efflux (Remm et al., 2022; Johnson et al., 2020). The use of efflux inhibitors for the treatment of tuberculosis, to complement and enhance existing treatment options has been discussed in numerous papers (Remm et al., 2022; Pule et al., 2016; Gupta et al., 2014; Laws et al., 2022; Szumowski et al., 2013). These studies have focussed upon verapamil, CCCP or plant natural products (Chen et al., 2018; Pule et al., 2016; Gupta et al., 2014), however, no efflux inhibitors are currently used clinically against tuberculosis (Pule et al., 2016). This is due to either a lack of safety data, for plant natural products, or toxic effects of inhibitors on eukaryotic systems, such as CCCP (Pule et al., 2016). Verapamil has a better safety profile, but causes serious adverse effects at higher concentrations (Pule et al., 2016). In contrast, vanoxerine has passed Phase I clinical trials without safety concerns arising in health volunteers (Obejero-Paz et al., 2015). Any adverse effects from vanoxerine were in patients with underlying structural heart disease and hence could be screened out during clinical trial testing and future use (Piccini et al., 2016).

The voltage sensitive dye DiOC_2_(3) is an indicator of membrane potential disruption (Chen et al., 2018; Chawla et al., 2012; Hudson et al., 2020; Li et al., 2019). Based on its use as a proxy, disruption of the electric potential (ΔΨ) is more likely to be a mechanism of action of vanoxerine. The ΔΨ disruption would cause the PMF of the cell to be dissipated, interfering with the energetics of the Mycobacterial cell, and hence leading to cell death (Chen et al., 2018; Feng et al., 2015). The dissipation of the PMF also would prevent the activity of efflux pumps, as many rely on the PMF to function (Remm et al., 2022), explaining the lack of ethidium bromide efflux. Vanoxerine is a cationic amphiphile at physiological pH, with a pK_a_ of 8.2 and a cLogP value of 5.3. Compounds with these properties have been shown to insert into lipid membranes and uncouple the PMF in bacterial inverted membrane vesicles, while having low mitotoxicity (Chen et al., 2018; Feng et al., 2015). In addition, membrane uncouplers have previously been reported to have several mechanisms of action (Feng et al., 2015) and this study does not preclude vanoxerine also having several mechanisms-of-action. The ability to interfere with cellular energetics has been shown to kill latent *M. tuberculosis* (Rao et al., 2008; Manjunatha et al., 2009), hence, vanoxerine should be tested for bactericidal effects in Mycobacteria during latency.

*M. tuberculosis* infections are always treated using a combination therapy, both to increase treatment efficacy and reduce levels of drug resistance development (Nahid et al., 2016; Zumla et al., 2013; Berry and Kon, 2009). Any new anti-mycobacterial drugs would be used within a combination therapy and thus, need to complement drugs within the current treatment regimen or novel drugs in development. Vanoxerine showed no interactions with the current or in development drugs clofazimine, FNDR-20081 or rifampicin, based on the FICs determined, although some shifts in the MIC_99_ were found (Kaur et al., 2021; Gopal et al., 2013). A reduction in MIC_99_ may allow lower doses of drugs to be used during treatment, hence, reducing associated side effects. Potential drug interactions may be masked by the concurrent upregulation of the MmpL5 efflux pump, suggested to efflux clofazimine and FNDR-20081, and inhibition of Mycobacterial efflux by vanoxerine (Hartkoorn et al., 2014; Andries et al., 2014; Remm et al., 2022; Kaur et al., 2021). Further testing is required to evaluate whether vanoxerine could potentiate the effects of mycolic acid or arabinogalactan biosynthesis inhibitors. In contrast, vanoxerine did not alter the MIC_99_ of bedaquiline, which may be due to these drugs having analogous mechanisms, both disrupting the PMF and hence having a similar cellular effect (Andries et al., 2005; Feng et al., 2015).

Comparing the RNA-sequencing data to other efflux inhibitors and uncouplers has given more perspectives. Phenothiazines have been shown to target the NADH dehydrogenase II, disrupting the electron transport chain, hence stopping efflux through PMF and ATP depletion (Remm et al., 2022). Transcriptomic data has shown phenothiazines to cause an increase in the transcript levels of *ndh, nuoE-G* and *icd1* (Dutta et al., 2010; Boshoff et al., 2004). Conversely, vanoxerine treatment did not affect the expression of *ndh* and *icd1*, while nuoE-G were all significantly down-regulated, suggesting NADH dehydrogenase II is not a target of vanoxerine. In contrast, the transcriptomic data showed a high degree of similarity to the 2-aminoimidazole class of compounds, which have been shown to dissipate the PMF and block the electron transport chain (Jeon et al., 2019, 2017). Treatment with both 2B8 (an 2-aminoimidazole) and vanoxerine resulted in upregulation of *mprA, sigB, sigE, mmpL5, mmpL8, mmpL10, rv3160c (bcg_3184c)* and *rv3161c (bcg_3185c)*, as responses to membrane stress, increasing membrane transporter/efflux and a putative dioxygenase and its regulator, respectively (Jeon et al., 2017). In addition, both compounds caused downregulation of both the mycolic acid (*fasI* and *fasII)* and peptidoglycan biosynthesis (*mur)* genes (Jeon et al., 2017). The main difference was vanoxerine did not induce transcription of the propionate detoxification genes, *prpC* and *prpD* or the *sigK* regulon (*sigK, rv0449c, mpt83, dipZ*), except for *mpt70*. Overall, the high similarity of the transcriptional responses of Mycobacteria to vanoxerine and 2B8, provides further evidence that vanoxerine is impacting the PMF and cellular energetics (Jeon et al., 2019, 2017).

Although the lipid analysis of *M. smegmatis* showed loss of TMM and TDM following vanoxerine treatment, the evidence suggests that vanoxerine does not directly target mycolic acid biosynthesis. This is supported by the fact that vanoxerine inhibits *C. glutamicum*, for which mycolic acids are not essential, with a comparable MIC_99_ to *M. tuberculosis*. Vanoxerine also had equal activity against *C. glutamicum* wild-type and the Δpks mutant. It is more likely that the PMF dissipation caused by vanoxerine has an indirect effect on the PMF-dependent transporter MmpL3, which transports TMM across the inner membrane (Su et al., 2019). This indirect inhibition is in contrast to the direct MmpL3 inhibition, which has been observed for many other antimycobacterial drugs (Li et al., 2019, p.3; Degiacomi et al., 2020, p.3). The observed reduction in the expression of the mycolic acid biosynthesis genes could also be an indirect result of MmpL3 inhibition leading to the accumulation of TMM and precursors in the cytoplasm. Loss of mycolates and downregulation of mycolic acid biosynthesis genes was also observed for the 2-aminoimidazole compounds, which are known to target the PMF (Jeon et al., 2017).

In summary, vanoxerine has been confirmed as an antimycobacterial drug with the ability to disrupt the membrane potential of Mycobacteria and should be included during pre-clinical testing of novel combination therapies. Future directions of research could include confirmation of electric potential disruption in *M. tuberculosis*, testing vanoxerine within a macrophage infection model, or against Mycobacteria in a hypoxia-induced latent state. In addition, analogues of vanoxerine could be synthesised to increase their potency against Mycobacteria and ideally reduce their effects on other human targets.

## Author Statements

### Authors and Contributors

AK designed the experiments. AK and AMJ performed the experiments. AK analysed the data. AK wrote the initial draft manuscript. AK, SB, and GS critically reviewed the manuscript.

### Conflict of Interest

The authors have no conflict of interest.

## Acknowledgments

We would like to thank Luke Alderwick for aiding preliminary work which later evolved into the current study. We would like to thank Tanya Parish and Jessica Blair for their suggestions of experimental work to undertake.

## Funding Information

This work was funded in part by the Wellcome Trust Doctoral Training Program Antimicrobials and Antimicrobial Resistance (grant reference: 108876/B/15/Z, to AK) and Microbiology Society (Harry Smith Vacation Studentship, to AMJ). For the purpose of open access, the author has applied a CC BY public copyright licence to any Author Accepted Manuscript version arising from this submission.

## Notes

https://www.ebi.ac.uk/ena/browser/view/PRJEB57729

### Competing Interest Statement

The authors have declared no competing interest.

