## Supplementary Material for "Repurposing Vanoxerine as a new antimycobacterial drug and its impact on the mycobacterial membrane"

### Supplementary Information:

**Supplementary Table 1: Table of Bacterial Strains used in the study**

| Microorganism | Strain | Plasmid/Mutant | Antibiotic Resistance | Reference |
|---|---|---|---|---|
| *Mycobacterium smegmatis* | $mc^2155$ | | | |
| *Mycobacterium smegmatis* | $mc^2155$ | pVV16 (empty) | Kanamycin | This study |
| *Mycobacterium smegmatis* | $mc^2155$ | pVV16-MtAroB | Kanamycin | This study |
| *Mycobacterium smegmatis* | $mc^2155$ | pTIC6a (empty) | Kanamycin | This study |
| *Mycobacterium smegmatis* | $mc^2155$ | pTIC6a-MtAroB | Kanamycin | This study |
| *Mycobacterium bovis* | BCG Pasteur | | | |
| *Mycobacterium bovis* | BCG Pasteur | ΔrecG | | (Batt et al., 2015) |
| *Escherichia coli* | BL21 (DE3) | pET28a-MtAroB | Kanamycin | This study |
| *Corynebacterium glutamicum* | 13032 | | | |
| *Corynebacterium glutamicum* | 13032 | Δpks | | (Gande et al., 2004) |
| *Enterococcus faecium* | 64/3 | | Rifampicin, fusidic acid | (Werner et al., 2003) |
| *Staphylococcus aureus* | SA01 | | | |
| *Klebsiella pneumoniae* | Ecl08 (KP02) | | | |
| *Acinetobacter baumanii* | AYE (AC05) | | | |
| *Pseudomonas aeruginosa* | PA14 | | | |

**Strain References:**


Batt, S.M., Cacho Izquierdo, M., Castro Pichel, J., et al. (2015) Whole Cell Target Engagement Identifies Novel Inhibitors of *Mycobacterium tuberculosis* Decaprenylphosphoryl-β- D -ribose Oxidase. *ACS Infectious Diseases*, 1 (12): 615–626. doi:10.1021/acsinfecdis.5b00065.

Gande, R., Gibson, K.J.C., Brown, A.K., et al. (2004) Acyl-CoA Carboxylases (accD2 and accD3), Together with a Unique Polyketide Synthase (Cg-pks), Are Key to Mycolic Acid Biosynthesis in Corynebacterianeae Such as Corynebacterium glutamicum and Mycobacterium tuberculosis. *Journal of Biological Chemistry*, 279 (43): 44847–44857. doi:10.1074/jbc.M408648200.

Werner, G., Willems, R.J.L., Hildebrandt, B., et al. (2003) Influence of Transferable Genetic Determinants on the Outcome of Typing Methods Commonly Used for Enterococcus faecium. *Journal of Clinical Microbiology*, 41 (4): 1499–1506. doi:10.1128/JCM.41.4.1499-1506.2003.

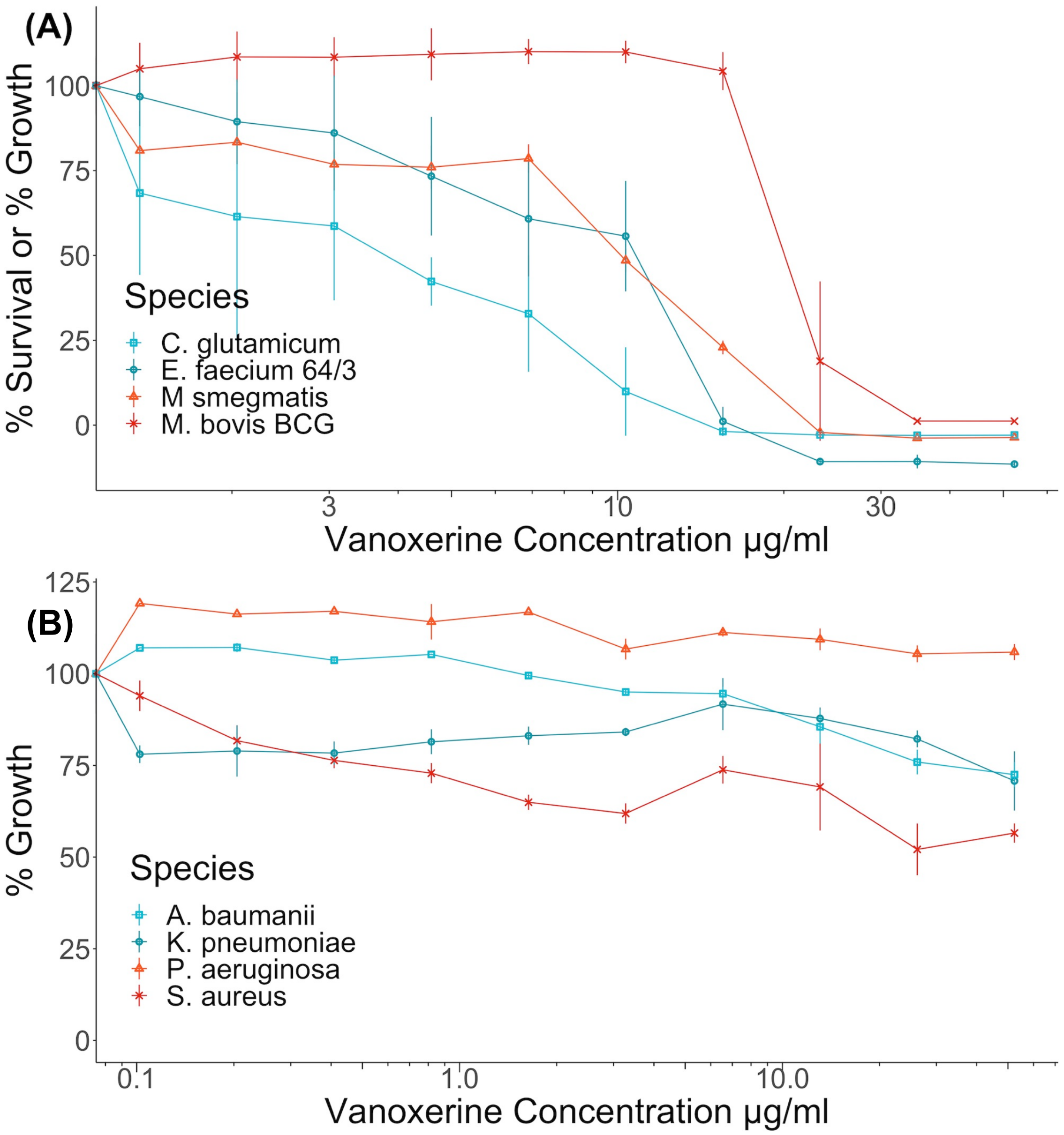


**Supplementary Figure 1: % Survival or % Growth curves of all tested species against Vanoxerine.** (A) Species vanoxerine could inhibit. (B) Species vanoxerine did not inhibit. Bacterial cultures were diluted to $OD_{600}$ = 0.05. A 2:3 serial dilution (A) or a 1:2 serial dilution (B) of vanoxerine was performed. The dilution series was transferred to an assay plate and the diluted cell cultures were added. The plates were incubated for 21 h (*M. smegmatis*), 23 h (*C. glutamicum*), 24 h (*E. faecium* 64/3*, A. baumanii, K. pneumoniae, P. aeruginosa & S. aureus*) or 6 days (*M. bovis* BCG). For *E. faecium* 64/3*, A. baumanii, K. pneumoniae, P. aeruginosa & S. aureus*, the $OD_{600}$ was measured. For *M. smegmatis, C. glutamicum* and *M. bovis* BCG, resazurin was added and the plates re-incubated for 1 h (*C. glutamicum*), 3 h (*M. smegmatis*) or 24 h (*M. bovis* BCG). Then the resorufin fluorescence was measured. In all cases, the % survival was calculated relative to a positive control (e.g., Rifampicin) and a negative control (DMSO only).

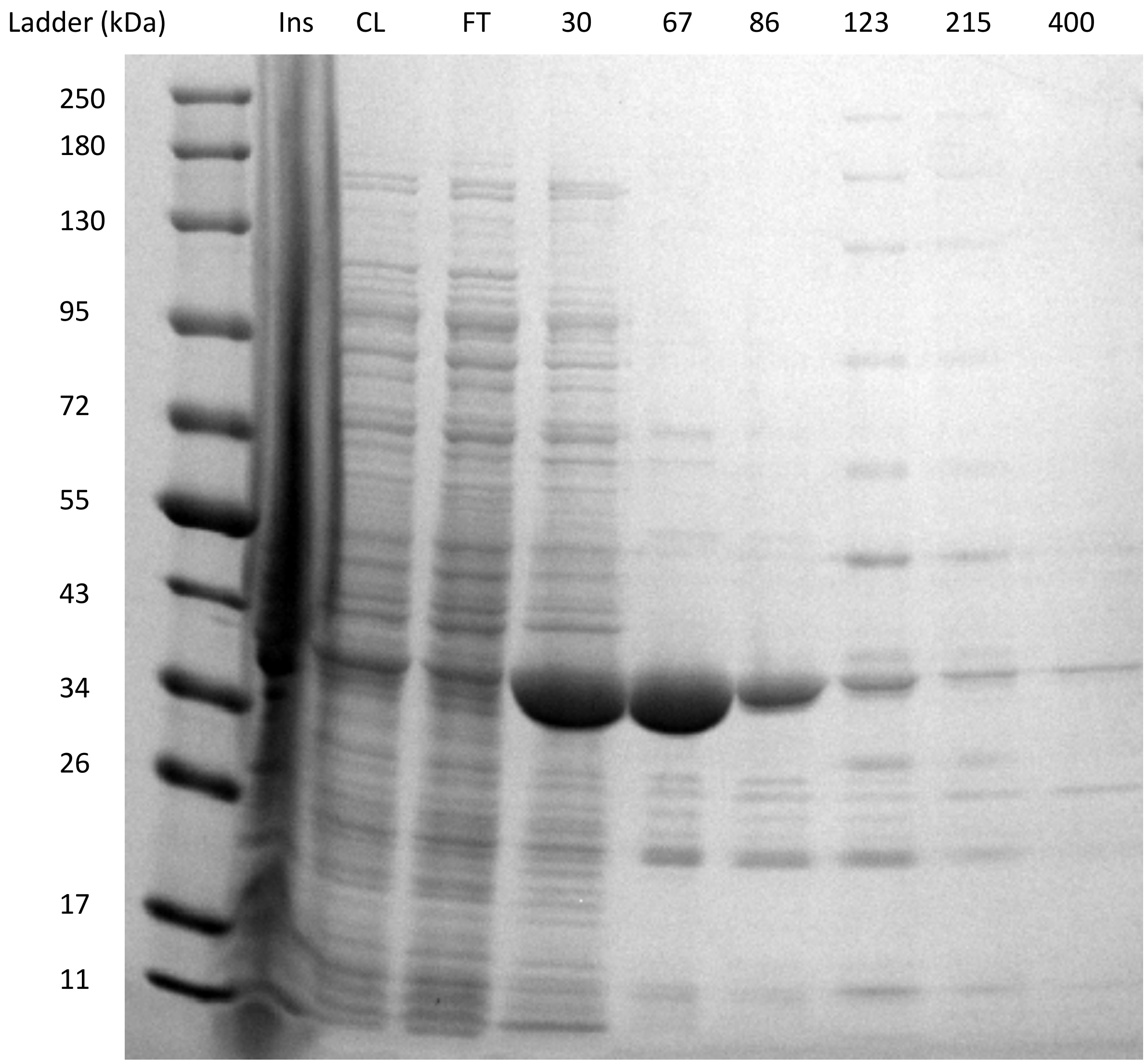


**Supplementary Figure 2: Mt-AroB (39 kDa) was successfully purified using a nickel NTA column.** *E. coli* BL21(DE3) cells containing the pET28a-MtAroB plasmid, had protein expression induced using IPTG for 18 hours. The cells were harvested, resuspended in buffer and lysed. This lysate was clarified, before being loaded onto a nickel NTA column. The column was washed with buffer containing 30 mM imidazole, before the protein was eluted, using increasing imidazole concentrations. This SDS-PAGE gel displays samples of each stage of purification, with Mt-AroB appearing at 39 kDa. Ins = insoluble fraction, CL = clarified lysate, FT = flow-through, 30 to 400 represent mM concentrations of imidazole in the buffer.

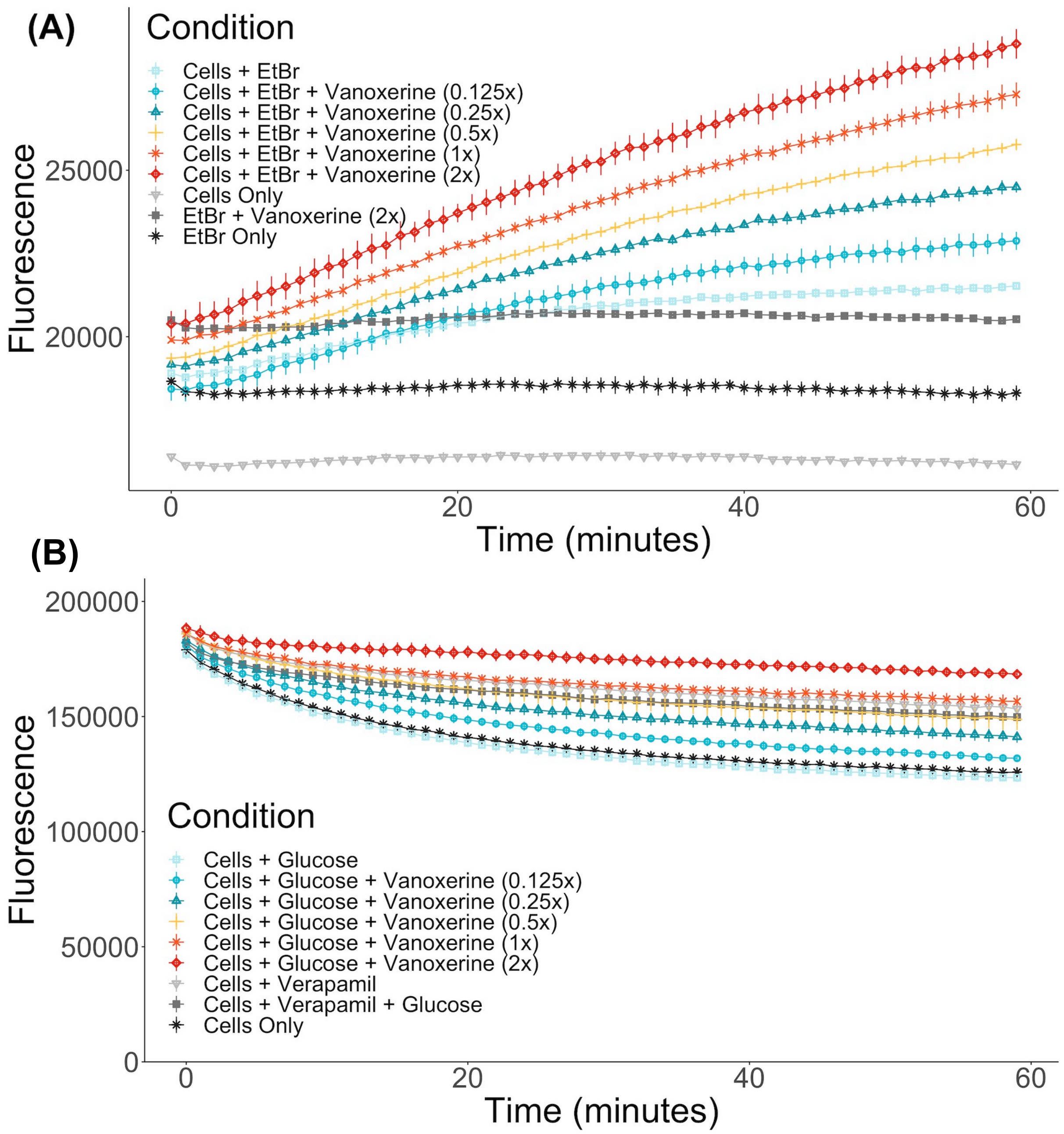


**Supplementary Figure 3: (A) Vanoxerine treated *M. bovis* BCG accumulated ethidium bromide at an increased rate.** *M. bovis* BCG was washed and re-suspended in PBS + 0.05% tween-80 + 0.4% glucose + 0.625 µg/ml ethidium bromide. The cells were immediately added to a 96-well plate containing varying vanoxerine concentrations and the fluorescence was measured for 1 hour. Vanoxerine $MIC_{99}$ = 26 µg/ml. N=3. **(B) *M. bovis* BCG efflux of ethidium bromide was inhibited by sub-inhibitory concentrations of vanoxerine.** *M. bovis* BCG was allowed to accumulate ethidium bromide (0.625 µg/ml) for 1 hour, using verapamil (50 µg/ml). Then the cells were resuspended in fresh PBS + 0.05% tween-80. Cells were either added directly or mixed with glucose (0.4%) before adding to a 96-well plate containing vanoxerine or verapamil. Vanoxerine $MIC_{99}$ = 26 µg/ml. N=3

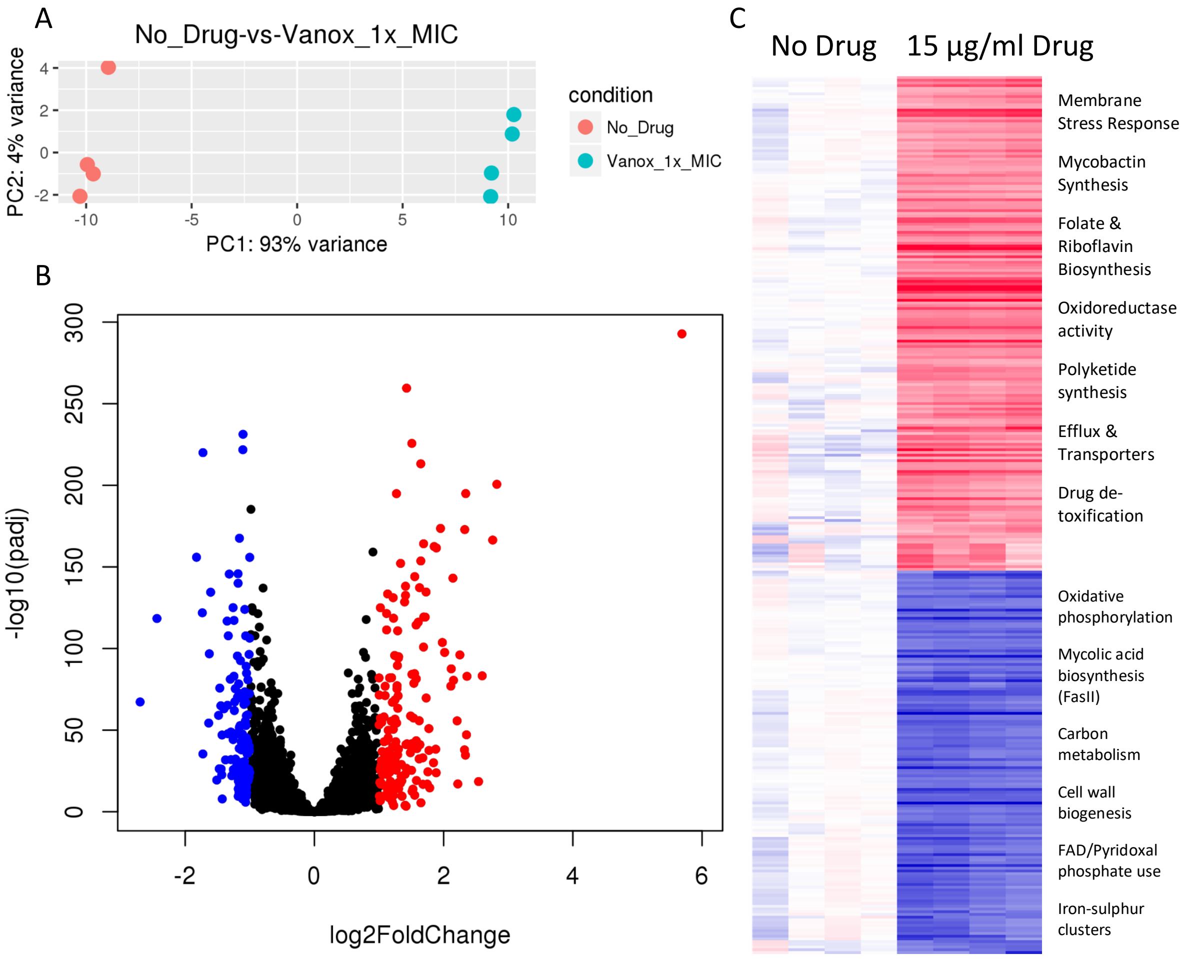


**Supplementary Figure 4: Transcriptomic changes following drug treatment of *M. bovis* BCG relative to a DMSO only control**. RNA sequencing was performed on four samples from each condition. (A) Principal component analysis highlighting the difference in transcriptional variation between DMSO-only controls and 15 µg/ml vanoxerine, following 8-hours of drug treatment. (B) Volcano plot indicating the significantly dysregulated genes. Highlighted points have a fold change > 2 and a p-adjusted value of <0.05; red = up-regulated, blue = down-regulated. (C) Clustered comparison of transcriptional differences between DMSO-only and 15 µg/ml vanoxerine. Biological functions of the dysregulated genes are highlighted.

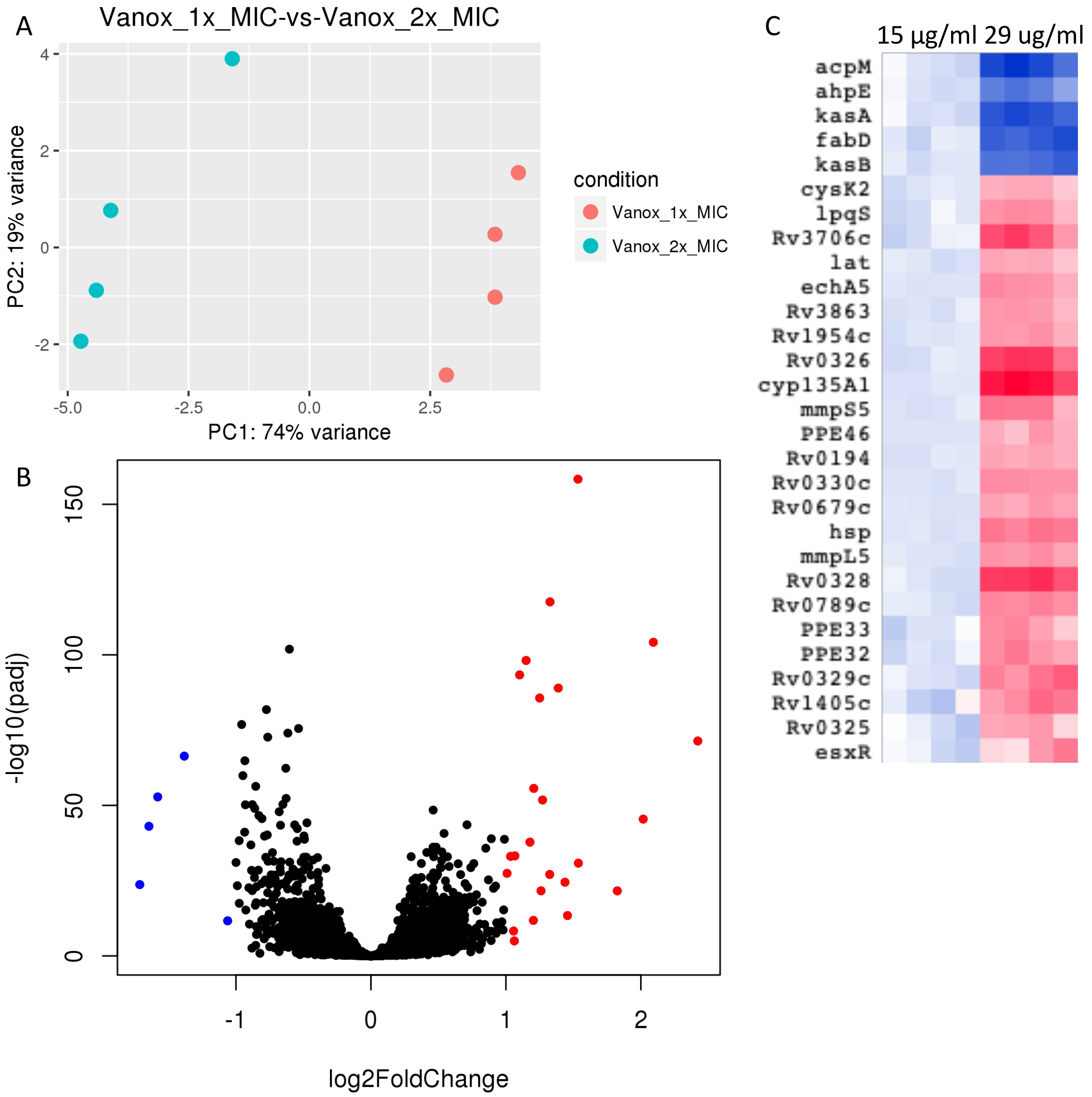


**Supplementary Figure 5: Transcriptomic changes following drug treatment of *M. bovis* BCG, comparing 15 to 30 μg/ml vanoxerine.** RNA sequencing was performed on four samples from each condition. (A) Principal component analysis highlighting the difference in transcriptional variation between 15 and 30 μg/ml of vanoxerine, following 8-hours of drug treatment. (B) Volcano plot indicating the significantly dysregulated genes. Highlighted points have a fold change > 2 and a p-adjusted value of <0.05; red = up-regulated, blue = down-regulated. (C) Clustered comparison of transcriptional differences between 15 and 30 μg/ml of vanoxerine. Dysregulated gene transcripts are named (equivalent TB naming is used if known).

**Supplementary Table 2: Significantly dysregulated genes comparing treatment of *M. bovis* BCG following treatment of 15 and 30 μg/ml of vanoxerine.** Data displayed in Supplementary Figure 6.

| ID | ID in TB | Functional Annotation | log2FoldChange | padj |
|---|---|---|---|---|
| *acpM* | *acpM* | Mycolic acid biosynthesis | -1.7106273 | 4.13E-24 |
| *kasA* | *kasA* | Mycolic acid biosynthesis | -1.6430428 | 4.24E-43 |
| *fabD* | *fabD* | Mycolic acid biosynthesis | -1.5780505 | 6.23E-54 |
| *kasB* | *kasB* | Mycolic acid biosynthesis | -1.3809422 | 6.23E-68 |
| *BCG_2265* | *Unknown* | Unknown | -1.118137 | 3.46E-11 |
| *ahpE* | *aphE* | Peroxidase | -1.0634044 | 4.20E-12 |
| *cysK2* | *cysK2* | Cysteine synthesis | 1.01017264 | 1.15E-28 |
| *lat* | *lat* | Lysine - glutamate conversion | 1.04721741 | 6.53E-28 |
| *PPE47* | *PPE47/PPE48 (Rv3021c)* | Unknown | 1.05591478 | 1.01E-13 |
| *esxR* | *esxR* | Secreted protein | 1.06344389 | 1.09E-05 |
| *PPE46* | *PPE46 (Rv3018c)* | Unknown | 1.06624228 | 1.89E-44 |
| *BCG_0231* | *Rv0194* | Multidrug efflux ATP-binding/permease protein | 1.10345536 | 4.80E-96 |
| *BCG_0728c* | *Rv0679c* | Unknown | 1.15139373 | 5.48E-98 |
| *BCG_3926* | *Rv3863* | Unknown | 1.17968576 | 3.14E-39 |
| *IS1606'* | *Rv0850* | Transposase | 1.19977257 | 6.15E-06 |
| *BCG_1993c* | *Unknown* | Unknown (DUF402) | 1.22347787 | 4.71E-59 |
| *mmpL5* | *mmpL5* | Efflux | 1.25195902 | 5.07E-87 |
| *lpqS* | *lpqS* | Lipoprotein of unknown function | 1.26228081 | 1.46E-22 |
| *echA5* | *echA5* | Fatty acid oxidation | 1.27325471 | 2.22E-53 |
| *PPE32* | *PPE32* | Unknown | 1.32645124 | 4.02E-28 |
| *BCG_0369c* | *eccE3 (Rv0292)* | ESX-3 secretion system protein EccE | 1.32867714 | 4.13E-120 |
| *PPE33a* | *PPE33a* | Unknown | 1.35099382 | 2.60E-11 |
| *BCG_0842c* | *Rv0789c* | Unknown | 1.3900481 | 6.68E-89 |
| *mmpS5* | *mmpS5* | Efflux | 1.4394337 | 1.95E-25 |
| *BCG_1466c* | *Rv1405* | Putative methyltransferase | 1.45830259 | 3.67E-14 |
| *hsp* | *hsp* | Stress response | 1.53579295 | 4.00E-158 |
| *BCG_0368c* | *Rv0329c* | Putative methyltransferase | 1.53961684 | 2.22E-31 |
| *BCG_3766c* | *Rv3706* | Unknown | 1.82697662 | 1.61E-22 |
| *BCG_0365* | *Rv0325* | Putative methyltransferase | 1.84468403 | 1.96E-44 |
| *BCG_0367* | *Rv0328* | Possible transcriptional regulator | 2.09420983 | 1.04E-105 |
| *cyp135A1* | *cyp135A1* | Cytochrome P450 | 2.42351955 | 7.71E-73 |

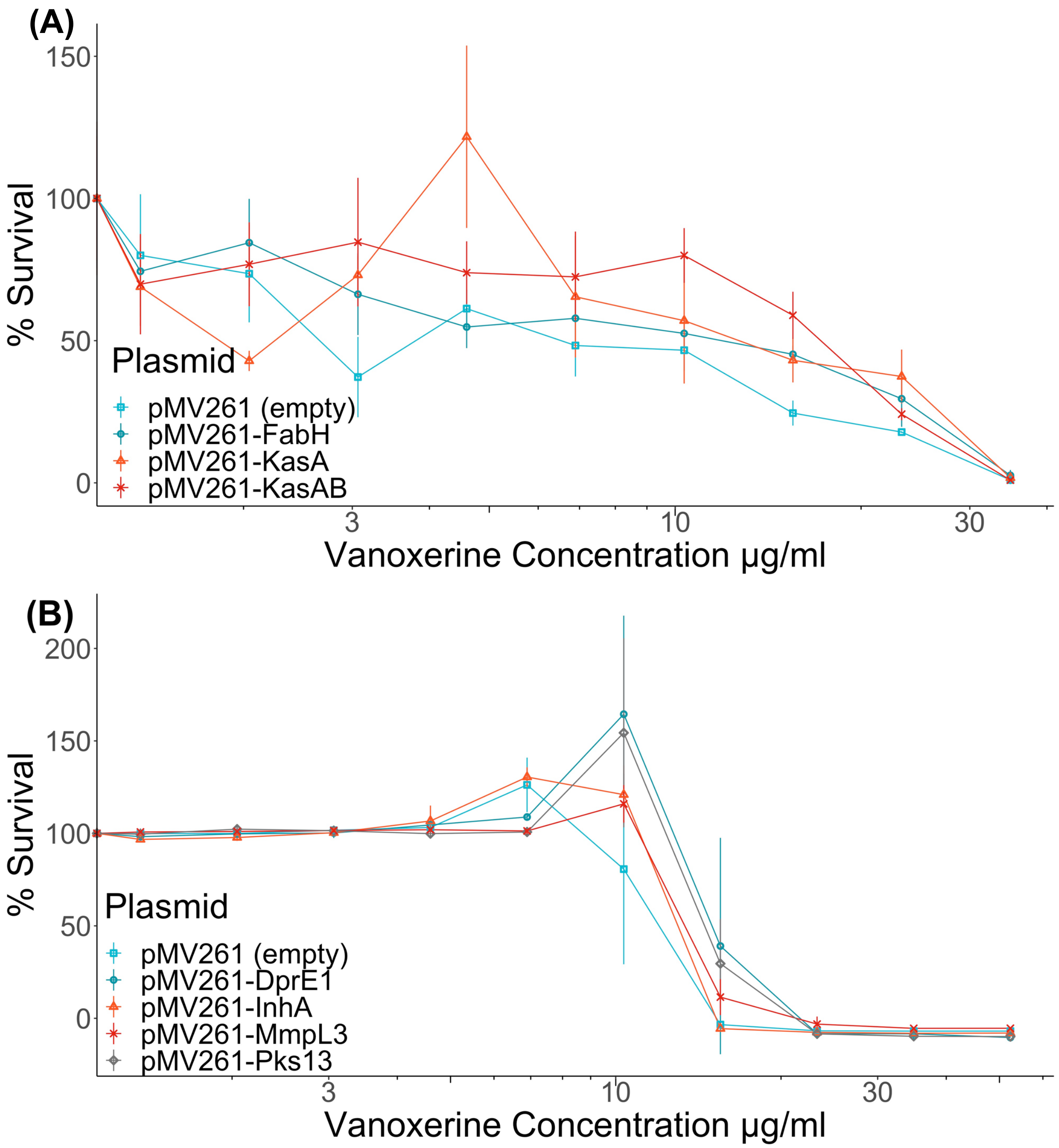


**Supplementary Figure 6: Mycobacteria over-expressing genes from the mycolic acid biosynthetic pathway had no impact on vanoxerine resistance.** (A) *M. smegmatis* (B) *M. bovis* BCG. Bacterial cultures were diluted to $OD_{600}$ = 0.05. A 2:3 serial dilution of vanoxerine was performed into DMSO, transferred to an assay plate (1 µl per well) and the diluted cell cultures was added (99 µl). The plates were incubated at 37 ˚C, for 21 h (*M. smegmatis*) or 6 days (*M. bovis* BCG). For *M. smegmatis* and *M. bovis* BCG, resazurin was added and the plates re-incubated for 3 h (*M. smegmatis*) or 24 h (*M. bovis* BCG). Then the resorufin fluorescence was measure. The %survival was calculated relative to a positive control and a negative control.
